# An Inflammatory Clock Predicts Multi-morbidity, Immunosenescence and Cardiovascular Aging in Humans

**DOI:** 10.1101/840363

**Authors:** Nazish Sayed, Tianxiang Gao, Robert Tibshirani, Trevor Hastie, Lu Cui, Tatiana Kuznetsova, Yael Rosenberg-Hasson, Rita Ostan, Daniela Monti, Benoit Lehallier, Shai Shen-Orr, Holden T. Maecker, Cornelia L. Dekker, Tony Wyss-Coray, Claudio Franceschi, Vladimir Jojic, François Haddad, José G. Montoya, Joseph C. Wu, David Furman

## Abstract

While many diseases of aging have been linked to the immunological system, immune metrics with which to identify the most at-risk individuals are lacking. Here, we studied the blood immunome of 1001 individuals age 8-96 and derived an inflammatory clock of aging (iAge), which tracked with multi-morbidity and immunosenescence. In centenarians, iAge was on average, 40 years lower than their corresponding chronological age. The strongest contributor to this metric was the chemokine CXCL9, which was involved in cardiac aging, affected vascular function, and down-regulated Sirtuin-3, a longevity-associated molecule. Thus, our results identify an important link between inflammatory molecules and pathways known to govern lifespan.

## Introduction

The role of the immune system in the maintenance of human health and protection against infections has been recognized for over a hundred years. However, it was only in the past few decades that it has become apparent that inflammatory components of the immune system are often chronically elevated in aged individuals and associated with an increased incidence of cancer, cardiovascular disease and neurodegenerative disorders, among others (Crusz and Balkwill, 2015; Kotas and Medzhitov, 2015; Liu et al., 2017). From these observations, it has been postulated that inflammation plays a critical role in regulating physiological aging (Franceschi and Campisi, 2014; Furman et al., 2017b). Furthermore, the well-established nine hallmarks of aging (Lopez-Otin et al., 2013); (1) genomic instability, (2) shortening telomere length, (3) epigenetic modifications, (4) loss of proteostasis, (5) deregulated nutrient sensing, (6) mitochondrial dysfunction, (7) cellular senescence, (8) stem cell exhaustion, and (9) altered intracellular communication, have all been shown to be caused, at least in part, by sustained systemic inflammation (Cavadas et al., 2016; Efeyan et al., 2015; Grivennikov et al., 2010; Hunter et al., 2007; Jurk et al., 2014; Lasry and Ben-Neriah, 2015; Nathan and Cunningham-Bussel, 2013; Oh et al., 2014; Thevaranjan et al., 2017; Alpert et al., 2019).

Contrary to the acute response, which is typically triggered by infection, chronic and systemic inflammation is thought to be triggered by physical, chemical or metabolic noxious stimuli (“sterile” agents) such as those released by damaged cells and environmental insults, generally termed “damage-associated molecular patterns” (DAMPs). This type of inflammation is associated with aging and characterized by being low-grade and persistent, ultimately leading to collateral damage to tissues and organs (Goldberg and Dixit, 2015; Kotas and Medzhitov, 2015). Despite the importance of this type of inflammatory reaction, there are currently no standard biomarkers to define a chronic inflammatory state and studies have generally yielded conflicting results (Franceschi et al., 2017; Morrisette-Thomas et al., 2014).

To better define biological markers of chronic inflammation and disease, we set out to establish a broad survey of immunity in over 1000 individuals (the Stanford 1000 Immunomes Project or Stanford 1KIP), wherein biological samples from 1001 subjects were obtained in the years 2007-2016 and comprehensively measured in a single facility; the Stanford Human Immune Monitoring Center (HIMC). At this center peripheral blood specimens were processed and analyzed using multiple technological platforms such as serum cytokines, cell subset composition, the cellular responses to multiple stimuli and the seropositivity to cytomegalovirus infection. In 902 subjects, a comprehensive health assessment using a 53-feature clinical questionnaire was also obtained.

Given the well-established importance of chronic inflammation for many human diseases and the lack of standard measures of it, we used deep learning methods to construct a metric for age-related chronic inflammation (iAge), which correlated with multiple morbidities and markers of immunosenescence, and was significantly lower in centenarians. In a validation study of 97 healthy older adults who were also monitored at the HIMC (Stanford) for the same inflammatory markers, along with a 27-feature cardiovascular phenotyping screening at the Stanford Cardiovascular Institute, we found that the most robust contributor to iAge, the interferon-related chemokine CXCL9, tracked with subclinical cardiac remodeling and arterial stiffness. This chemokine was also expressed in large quantities by aged endothelium derived from human induced pluripotent stem cells (hiPSC), suppressed vascular function in mice aortic tissue, and inhibited expression of the cardio-protective longevity-associated deacetylase Sirtuin-3.

Thus, our results define healthy versus unhealthy aging and identify a link between inflammatory molecules of the immune system and known pathways governing lifespan in humans and in model organisms.

## Results

### Study design of the Stanford 1000 Immunomes Project

During the years 2007 to 2016, blood samples from ambulatory subjects (*N* = 1001) (339 males and 662 females) from age 8 to >89 (Fig. S1 and Fig. S2) who had been recruited at Stanford University (the Stanford 1000 Immunomes Project, or Stanford 1KIP) for a longitudinal study of aging and vaccination (Blazkova et al., 2017; Brodin et al., 2015; Furman et al., 2017b; Furman et al., 2014; Furman et al., 2013; Furman et al., 2015; Price et al., 2013; Roskin et al., 2015; Shen-Orr et al., 2016; Wang et al., 2014), and for an independent study of chronic fatigue syndrome (Montoya et al., 2017), from which only healthy controls were included. Inclusion and exclusion criteria can be found under the Methods section. For all samples of the Stanford 1KIP, deep immune phenotyping was conducted at the Stanford Human Immune Monitoring Center (HIMC), where peripheral blood specimens were processed and analyzed using rigorously standardized procedures (Maecker et al., 2012). Serum samples were obtained and used for protein content determination (including a total of 50 cytokines, chemokines and growth factors) (*N* = 1001) and to assess cytomegalovirus status (*N* = 748), a major determinant of immune system variation (Brodin et al., 2015; Furman et al., 2015). Peripheral blood mononuclear cells or whole blood samples were used for the determination of cellular phenotypes and frequencies (*N* = 935), and for investigation of *in vitro* cellular responses to a variety of cytokine stimulations (*N* = 818). Extended clinical report forms were collected from 902/1001 subjects, of which 299 were males and 603, females (Table S1 and S2). A total of 37 additional older adults (19 centenarians and 18 controls) were included and screened for serum protein content to derive iAge on these extremely long-lived humans.

### Deep learning analysis of circulating immune cytokines and chemokines to derive a signature for age-related chronic inflammation

Given the increasingly recognized effect of low-grade systemic chronic inflammation in the development of a wide variety of diseases associated with aging, especially in cardiovascular disease (Furman et al., 2017b; Ridker et al., 2017), we set out to construct a metric for age-related chronic inflammation that could summarize an individual’s inflammatory burden. This type of inflammation is thought to ensue as a maladaptive reaction in response to exposure to tissue damage, metabolic dysfunction and environmental insults (collectively referred to as the “exposome”) (Goldberg and Dixit, 2015; Kotas and Medzhitov, 2015). In contrast to the acute inflammatory response, for which a number of secreted molecules (such as C-reactive protein, IL-1β, IL-6 and TNF-α) have been validated; in age-related chronic inflammation, no standard cytokine signature exists (Franceschi et al., 2017; Morrisette-Thomas et al., 2014). Thus, we undertook an unbiased approach to compactly represent the non-linear structure of the cytokine network. To do so, we used a deep learning method called guided auto-encoder (GAE) to circulating immune protein data encompassing a total of 50 cytokines, chemokines and growth factors from all 1001 subjects.

In the GAE method, the original data are combined *in silico* into a small number of ‘codes’ by a non-linear function. In this initial step, a first ‘hidden layer’ of a neural network is built. By combining each ‘code’ from the hidden layer into a new set of codes, iteratively, a second and third hidden layers are generated (for example, see Methods). This process aims at eliminating the noise and redundancy in the data, yet retaining the most relevant information, such that a robust GAE model is able to accurately predict from the last set of codes (last hidden layer), the original raw data (data reconstruction). In our analysis, the computational task was to predict the original cytokine data and use age as a second output variable, such that the predicted values represent a given individuals’ age-related chronic inflammation level, or inflammatory clock.

To test the robustness and quality of the GAE method, we compared the accuracy of the age prediction against other widely used dimensionality reduction methods that use linear equations, such as the elastic net and principal component analysis (PCA), as well as those involving non-linear equations, such as plain auto-encoders and neural networks (Fig. S3a and b). Overall, the GAE method outperformed other methods in predicting chronological age (*P* < 0.05) with the exception of the comparison with a plain neural network (*P* = 0.54) (Fig. S3b). These results indicate that the phenomenon of low-grade chronic inflammation in aging humans is best modeled using non-linear methods, and based on these, one can derive a metric for chronic inflammation that accurately predicts chronological age in the population, while preserving the biological information related to the total inflammatory burden as measured by the level of circulating immune proteins.

### The inflammatory clock (iAge) is correlated with multi-morbidity

To gain further insights into the effect of age-related chronic inflammation on age-related pathology, we computed a regression analysis between the sum of age-related disease (multi-morbidity) and iAge. The items analyzed included different diseases and physiological systems: cancer, cardiovascular, respiratory, gastrointestinal, urologic, neurologic, endocrine-metabolic, musculoskeletal, genital-reproductive and psychiatric. All these disease features were binary.

For these analyses, we controlled for age, BMI, sex, CMV and high cholesterol (also a binary category) (Fig. 1a), given the reported effect of each of these variables in the etiology of age-related pathologies. We observed a significant correlation between iAge and multi-morbidity in the older adults in this study (>60 years old) (*N* = 285, *P* < 0.0001) (Fig. 1b), but not with any individual disease item (Table S3). In addition to this analysis, we envisioned an unbiased approach to select predictors of multi-morbidity based on the available data for all 902 subjects while controlling for the age effect. To do so, we used a shrinkage method for variable selection by cross-validation, called the Elastic Net, which has been increasingly used in immunology, aging and other medical fields over the past years. We applied differential penalties for each potential predictor such that the machine learning procedure would ‘force’ age to be selected, while imposing a stringent penalty to all other features, so that the variables selected do not correlate with age (Fig. 1c). The Mean Absolute Error (MAE) for the prediction of multi-morbidity was 0.41 (Fig. 1d) with eighteen selected features, including iAge, high cholesterol and BMI (Fig. 1e), and immune parameters such as total CD8 (+) T cells, plasmablasts and transitional B cells (negative predictors) and IgD+CD27- and IgD-CD27-B cells, effector CD8 (+) T cells, total lymphocytes and monocytes, and central memory T cells (positive predictors) (Fig. 1f).

**Figure 1.**
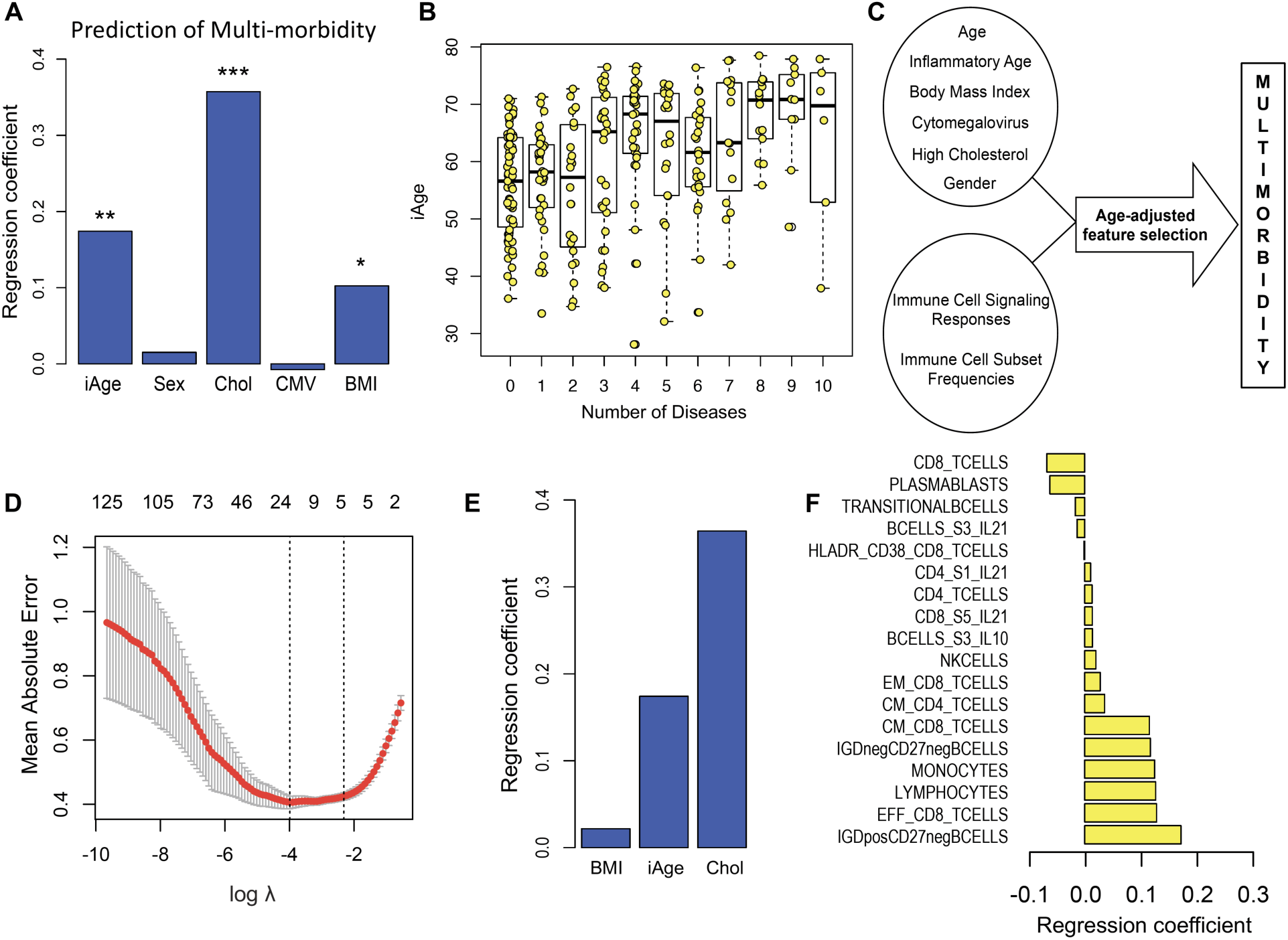
Inflammatory age predicts multi-morbidity. Ten age-related disease items were selected to characterize the clinical significance of inflammatory age derived from the analysis of 50 cytokines (*N* = 1001), using the guided autoencoder method. The items analyzed included different diseases and physiological systems: cancer, cardiovascular, respiratory, gastrointestinal, urologic, neurologic, endocrine-metabolic, musculoskeletal, genital-reproductive and psychiatric. All these disease features were binary. A significant regression coefficient was obtained for inflammatory age in the prediction of sum of diseases (multi-morbidity), in the older population analyzed (>60 years old, *N* = 285). Covariates included age (not shown), BMI, sex, CMV and high cholesterol (also a binary category) (**A**). In (**B**), age-adjusted inflammatory age is plotted against sum of diseases. To unbiasedly select for predictors of comorbidity based on available data for all 902 subjects while controlling for the age effect, age-adjusted cross-validation was performed (**C**). By applying differential penalty values for each regressor, age variable is ‘forced in’, while imposing a stringent penalty (the lasso penalty) to all other features, so that selected variables do not correlate with age. A Mean Absolute Error (MAE) for the prediction of comorbidity of 0.41 is observed (**D**). Eighteen features are selected including inflammatory age, high cholesterol and BMI (**E**) and immune parameters such as total CD8 (+) T cells, plasmablasts and transitional B cells (negative predictors) and IgD+CD27- and IgD-CD27-B cells, effector CD8 (+) T cells, total lymphocytes and monocytes, and central memory T cells (positive predictors) (**F**).

Collectively, these results show that the inflammatory clock is a metric for overall health linked to multiple diseases associated with aging.

### The inflammatory clock (iAge) is correlated with immunosenescence

Canonical acute inflammation proteins (C-reactive protein, Interleukin-6, etc.) have been associated with immunosenescence in previous studies, but the relationship with systemic chronic inflammation has not yet been established. To investigate this link, we first used a well-known marker for immunosenescence (the frequency of naïve CD8 (+) T cells) and estimated the contribution of iAge after controlling for Age, CMV, and sex by a multiple regression model. Not surprisingly age was the strongest contributor to changes in naïve CD8 (+) T cells followed by iAge, CMV (negative contributors) and sex (frequency of total CD8 (+) T cells in females was 24% vs. 30% in males) (Fig. 2a).

**Figure 2.**
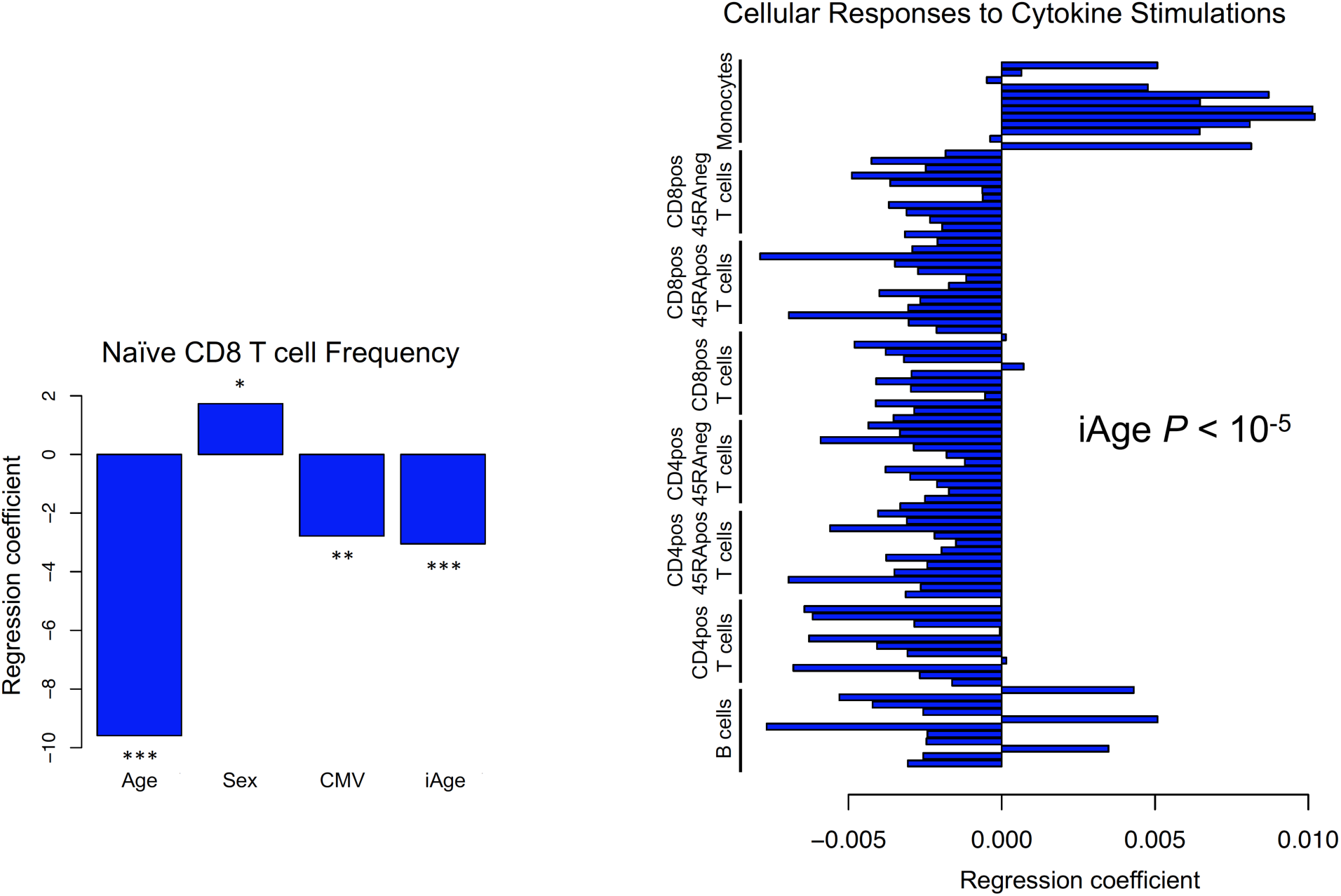
Inflammatory age correlates with immunosenescence. A well-known marker for immunosenescence (naïve CD8 (+) T cells) was used to examine the potential contribution of iAge to this condition. In a multiple regression model, iAge was significantly correlated with the frequency of naïve CD8 (+) T cells to a similar extent to CMV positivity. Chronological age was the strongest contributor (*P* < 10^-15^), followed by iAge (*P* < 10^-5^), CMV (*P* < 10^-3^) and gender (*P* = 0.012) (A). The activation of multiple pathways was measured using the phospho-flow method (see Method) in B cells, CD4 (+) T cells (and the CD45RA (+) and CD45RA (-) subsets), CD8 (+) T cells (and the CD45RA (+) and CD45RA (-) subsets) and in monocytes. iAge is consistently negatively correlated with B cell and T cell responses to cytokine stimuli, and positively correlated with monocyte responses (B) (*P* < 10^-5^ by self-contained test of modified Fisher’s combined probability (Dai et al., 2014)).

To examine the effect of chronic inflammation in the immune response, we used a functional immune assay (phospho-flow) in which cells are stimulated *ex vivo* and the phosphorylation of various intracellular proteins is measured by using antibodies against phosphorylated forms of these proteins. In particular, we selected the responses to four independent stimuli (Interferon-alpha, Interleukin-6, Interleukin-10 and Interleukin-21) which were consistently measured in a total of 818 individuals and measured the fold-increase in phospho-STAT1, -STAT3 and - STAT5 in B cells, total CD4 (+) T cells (and the CD45RA(+) and CD45RA(-) subsets), total CD8 (+) T cells (and the CD45RA(+) and CD45RA(-) subsets), and in monocytes, totaling 96 conditions. We conducted multiple regression analysis controlling age, CMV and sex. Strikingly, there was a general decrease of the B cell and T cell responses to stimuli and an overall potentiation of the monocyte responses associated with increasing iAge (combined *P* < 10^-5^) (Fig. 2b).

These results demonstrate that iAge is an important immune predictor of immune function decline (immunosenescence) and can be used as a ‘metric’ for immunological health.

### CXCL9 is an important component of the inflammatory clock and correlates with cardiovascular aging in healthy adults

Given the importance of chronic inflammation in cardiovascular disease and mortality (Furman et al., 2017a; Proctor et al., 2015), we explored potential mechanisms by which the inflammatory clock relates to cardiovascular dysfunction. To do so, we first aimed at analyzing which factors contribute the most to iAge. We computed the most variable jacobians (first order partial derivative of iAge) to decomposing the inflammatory clock into single immune protein features. We found both positive and negative contributors to iAge (Fig. 3a). The top 15 most variable jacobians were CXCL9, EOTAXIN, Mip-1α, LEPTIN, IL-1β, IL-5, IFN-α and IL-4 (positive contributors) and TRAIL, IFN-γ, CXCL1, IL-2, TGF-α, PAI-1 and LIF (negative contributors) (Fig. 3a).

**Figure 3.**
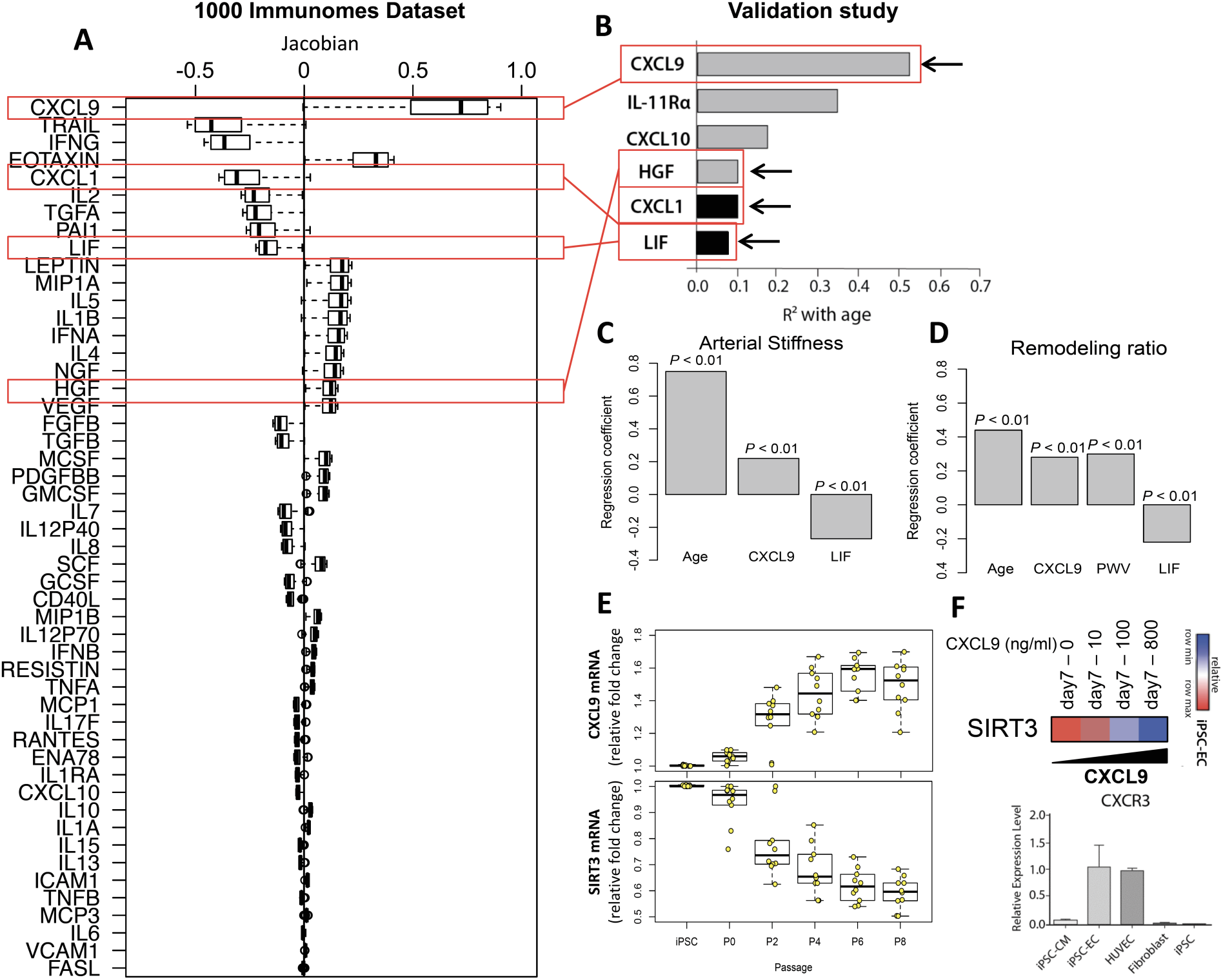
Endothelial cells derived from hiPSCs produce CXCL9, which predicts cardiovascular aging in healthy subjects independently of age. Decomposition of the inflammatory score was conducted by estimating the most variable jacobians (first order partial derivative of inflammatory age) (**A**). Both positive and negative contributors to inflammatory age are observed. The top 15 most variable jacobians are CXCL9, EOTAXIN, Mip-1α, LEPTIN, IL-1β, IL-5, IFN-α and IL-4 (positive contributors) and TRAIL, IFN-β, CXCL1, IL-2, TGF-α, PAI-1 and LIF (negative contributors) (A). In a validation study, 97 healthy adults (age 25-90) well matched for cardiovascular risk factors, were selected form a total of 151 recruited subjects. Immune protein analysis was conducted in samples from these subjects. CXCL9, HGF, CXCL1 and LIF were found to change in the same direction in both the Stanford 1KIP and the validation cohort (**A and B**). Cardiovascular age was estimated using 3 parameters: (1) aortic pulse wave velocity, a measure of vascular stiffness; (2) relative wall thickness (RWT), a measure of ventricular remodeling, and (3) early diastolic mitral annular velocities (e’), a measure of ventricular relaxation. After adjusting for age, sex, BMI, heart rate, systolic blood pressure, fasting glucose and total cholesterol to HDL ratio, positive correlations were obtained between CXCL9 and PWV (*R* = 0.22) and RWT (*R* = 0.3) (*P* < 0.05), and negative correlations were observed between LIF and PWV (*R* = −0.27), and RWT (*R* = −0.22) (**C and D**). No variable included in the models had high co-linearity as suggested by variance inflation factors (VIF) < 3 for each factor. Induced pluripotent stem cells (hiPSCs) were obtained from isolated fibroblasts (*N* = 5, in duplicates) using the Yamanaka factors (Takahashi and Yamanaka, 2006) and differentiated them into endothelial cells (hiPSC-ECs) under well-defined conditions as previously described (Hu et al., 2016). Expression levels of CXCL9 and SIRT3 were measured by RT-PCR as described under Methods. A significant age-dependent increase in CXCL9 mRNA expression levels is observed (*P* < 0.01), which reaches a plateau after the sixth cell passage (**E**). Concomitant with the increase in CXCL9, down-regulation in SIRT3 mRNA can be observed after the second cell passage (*P* < 0.01) (E). Addition of increasing doses (10 to 800 ng/ml) of exogenous CXCL9 to young (day 7) hiPSC-ECs induces down-regulation of SIRT3 mRNA expression (F). Expression of the CXCL9 receptor, CXCR3, was measured in young cardiomyocytes derived from hiPSCs (hiPSC-CM) as well as in hiPSC-ECs, HUVEC cells, freshly isolated fibroblasts and hiPSCs. Elevated expression is observed in hiPSC-ECs, HUVEC cells but not in other cell types (**F**) suggesting that the endothelium but no other cell subsets is target of CXCL9 and potentially other CXCR3 ligands as well.

To validate our findings, we conducted a follow-up study in a group of 97 healthy adults (age 25-90 years old) who were selected form a total of 151 recruited subjects, using strict selection criteria, and who were well matched for cardiovascular risk profiles (see Methods) (Table S4). Inflammation markers were measured using a 48-plex panel, and aortic pulse wave velocity (PWV) and echocardiograms were obtained from all the study participants. RWT was calculated as stated previously (see Methods).

In this healthy cohort, from the 48 circulating immune proteins, only 6 were significantly correlated with age (*P* < 0.05), and 4 were also found in the iAge decomposition analysis of the Stanford 1KIP dataset (Fig. 3b). Among them CXCL9, IL-11Rα, CXCL10 and HGF increased with aging, while CXCL1 and LIF, decreased (Fig. 3b). iAge predicted pulse-wave velocity after controlling for age and other confounders (*P*<0.001). To identify single inflammatory markers associated with cardiovascular aging that were independent of age and potentially present in healthy individuals, we next performed multiple regression hierarchical analysis on selected inflammatory markers associated with aging in this cohort (CXCL9 and LIF) and the cardiovascular measurements obtained from all the individuals studied (see Methods). For these analyses, we controlled for age, sex, BMI, heart rate, systolic blood pressure, fasting glucose and total cholesterol to HDL ratio. At a *P*-value < 0.01, we found a positive correlation between CXCL9 and the cardiovascular aging markers PWV (*R* = 0.22), a measure of arterial stiffness, and RWT (*R* = 0.3), a measure of cardiac remodeling; and a negative correlation between LIF and PWV (*R* = −0.27), and RWT (*R* = −0.22) (Fig. 3c and d).

Taken together, these results show that subclinical cardiac tissue remodeling and increased arterial stiffness can be found in otherwise healthy individuals with elevated levels of CXCL9 levels and low levels of LIF.

### Aging endothelial cells express high levels of CXCL9, which induces mRNA down-regulation of the cardio-protective SIRT3

Long standing evidence have suggested a critical role for the endothelium in the etiology of hypertension and arterial stiffness, and more recent work has also shown that more advanced signs of cardiovascular aging such as tissue remodeling and cardiac hypertrophy are often preceded and may be initiated by malfunctioning of aged endothelia (Castellon and Bogdanova, 2016; Harvey et al., 2015; Kamo et al., 2015). In order to study the role of CXCL9 in this process, we induced human fibroblasts obtained from 5 donors to a pluripotent stem cell (hiPSC) phenotype by using the Yamanaka factors (Takahashi and Yamanaka, 2006), and subsequently differentiated them into endothelial cells (hiPSC-ECs). To model the effect of aging, hiPSC-ECs were passaged numerous times as indicated (Fig. 3e), under well-defined conditions as previously described (Hu et al., 2016), and the mRNA expression levels of CXCL9 and SIRT3 were measured. Sirtuin-3 is an important deacetylase with cardio-protective properties involved in mitochondrial homeostasis, stem cell and tissue maintenance during aging, and linked to the beneficial effects of diet, caloric restriction and exercise in maintaining cardiovascular health and longevity (Bonkowski and Sinclair, 2016; Lu et al., 2016). We observed a time-dependent increase in CXCL9 transcript levels, which was concomitant with a drop in SIRT3 expression (Fig. 3e), and with the decrease in the number of vascular networks formed by the endothelial cells.

Treatment of young hiPSC-ECs (at day 7) with increasing doses of CXCL9 (10 to 800 ng/mL) also down-regulated SIRT3 expression (Fig. 3f), indicating that young endothelia is a target of CXCL9 from other sources, and can down-regulate SIRT3 expression upon exposure to CXCL9. In addition, we confirmed expression of the CXCL9 receptor, CXCR3 in young hiPSC-EC but not hiPSC-derived cardiomyocytes (Fig. 3f), indicating that CXCL9 can act both in a paracrine fashion, wherein increasing levels of this chemokine from immune sources affect endothelial cell function, and in an autocrine fashion on endothelial cells likely producing a positive feedback loop where increasing doses of CXCL9 and expression of its receptor in these cells leads to cumulative deterioration of endothelial function in aging.

To validate our findings that aging ECs express elevated levels of CXCL9, which in turn down-regulate SIRT3, we assessed the expression levels of both CXCL9 and SIRT3 in CXCL9 knockdown (KD) hiPSCs. Specifically, CXCL9 was KD in human hiPSCs using short hairpin RNA (shRNA) (expression was reduced by ∼75%, Fig. 4a) and then differentiated to ECs. These hiPSC-ECs were then serially cultured to passage 8 so as to mimic cellular aging with cells collected at every passage and RNA extracted for quantitative PCR. Consistent with our previous results (Fig. 3e-f), an increase in the passage number of hiPSC-ECs showed an increase in CXCL9 expression in scramble-treated hiPSC-ECs (Fig. 4b). However, in contrast, CXCL9-KD hiPSC-ECs failed to show an increase in CXCL9 expression as the hiPSC-ECs were passaged (Fig. 4b). Importantly, SIRT3 expression, which was down-regulated in a time-dependent manner in the aging hiPSC-ECs, showed a reversal in their levels in CXCL9-KD hiPSC-ECs (Fig. 4c).

**Figure 4.**
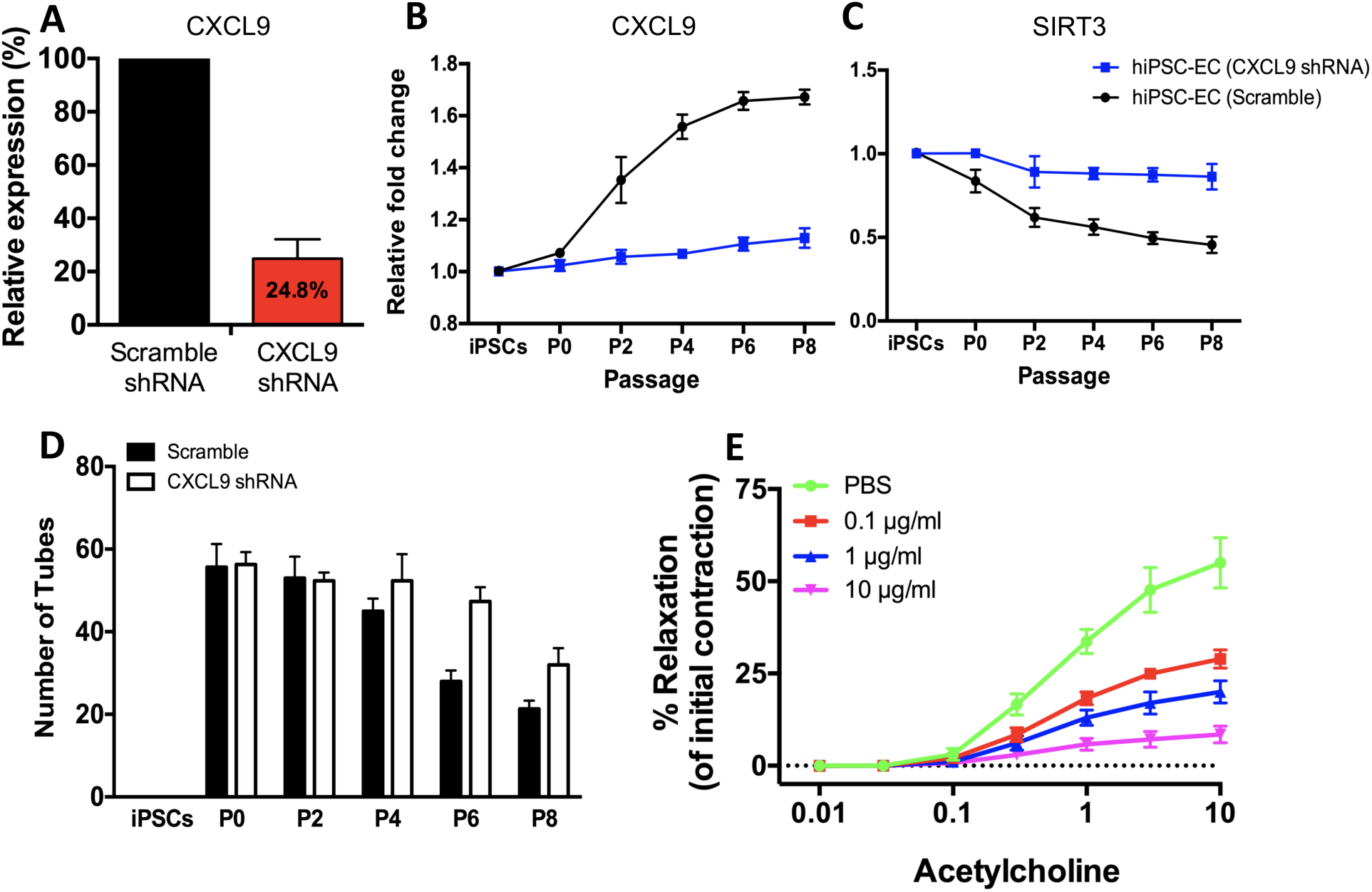
CXCL9 impairs endothelial function. To validate the association between CXCL9 expression and SIRT3 in hiPSC-ECs, CXCL9 was knockdown (KD) in hiPSCs using shRNA. Quantitative PCR data show significant knockdown of CXCL9 in hiPSCs following transfection with CXCL9-shRNA compared to scramble controls (**A**). hiPSC-ECs failed to show an increase in CXCL9 expression in CXCL9-KD cells as they were passaged when compared to scramble controls (**B**). On the contrary, CXCL9-KD hiPSC-ECs show reversal of their SIRT3 levels in serially passaged hiPSC-ECs when compared to scramble controls (**C**). Quantification of the number of capillary-like networks formed by scramble- and CXCL9-KD hiPSC-ECs show that CXCL9-KD hiPSC-ECs retain their capacity to form tubular networks even at later passages when compared to scramble controls that showed impaired tube formation at later passages of hiPSC-ECs (**D**). Line graph of percent relaxation of mouse thoracic aortic sections to Acetylcholine show impaired vascular reactivity to increasing concentrations of CXCL9, suggesting CXCL9 impairs vascular function (**E**). Significance of impaired vascular reactivity was determined by 2-way ANOVA, followed by a Bonferroni post-test, N=3, n=3-4, *p<0.05.

Next, we assessed the effects of CXCL9 on the endothelial phenotype by assessing the function of these hiPSC-ECs (scramble- and CXCL9-KD) at different passages (P0 to P8) for their capacity to form networks of tubular structures so as to mimic angiogenesis (Fig. S4). Consistent with our previous data we found that scramble-treated hiPSC-ECs formed networks of tubular structures at an earlier passage, however showed impaired tube formation at a later passage (Fig. 4d). In contrast, CXCL9-KD hiPSC-ECs retained their capacity to form tubular networks even at a later passage, suggesting that CXCL9 impairs endothelial function.

Last, we investigated the effect of CXCL9 on vascular function, by incubating mouse thoracic aortic sections with increasing concentrations of recombinant mouse CXCL9. The specimens were incubated with the prostaglandin agonist U46619 and concentration-dependent contraction curves were created. Subsequently, concentration-dependent relaxation curves of Acetylcholine were conducted on the vessels. As shown in Fig. 4e, a dose-dependent effect of CXCL9 is observed on vasorelaxation in treated aortas versus controls, demonstrating that CXCL9 impairs vascular function, possibly contributing to arterial stiffness and premature aging of the cardiovascular system.

### Centenarians have a lower inflammatory age compared to their chronological age

To investigate the relationship between inflammatory age and longevity we computed an inflammatory index (inflammatory age minus chronological age) in an additional cohort of 37 subjects, 18 of which were 50-79 years old and 19 centenarians, except for 1 who was 99 years old at the time of blood extraction. Despite a significant difference in the inflammatory index between centenarians and control older adults group (−36.5 vs −1.4, *P* = 10^-6^) (Fig. 5), a large variance was observed in the inflammatory index of centenarians (range −83.5, 12.1) suggesting that there may be other mechanisms apart from inflammation conferring disease protection and long lifespan to these subjects.

**Figure 5.**
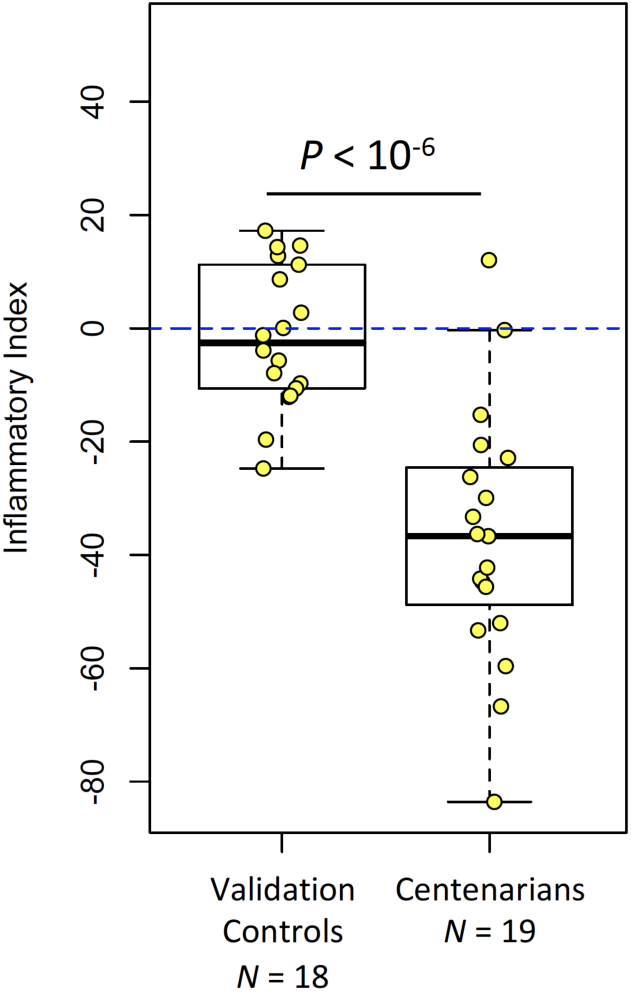
Centenarians have a low inflammatory index. Comparison of the inflammatory index (inflammatory age minus chronological age) was computed between a healthy group of older adults (*N* = 18, age range 50-79 years old) and centenarian subjects (*N* = 19, age range 99- 107). A significant difference in the inflammatory index is observed between centenarians and the control older adults’ group (−36.5 vs −1.4, *P* = 10^-6^) despite the large variance observed in the inflammatory index of centenarians (range −83.5, 12.1).

These results demonstrate that in long-lived humans iAge is significantly lower than the corresponding chronological age, suggesting that the inflammatory etiology of cardiovascular disease and multi-morbidity may translate into increased mortality.

## Discussion

In this study, we conducted extensive immune monitoring in a large cohort of 1001 subjects (the Stanford 1KIP) and used artificial intelligence methods to developed reference values for age-related chronic inflammation or “inflammatory age”, which correlated with multi-morbidity and immunosenescence. A major contributor to this inflammatory age metric, CXCL9, was validated as an indicator of cardiovascular pathology independent of age in an otherwise a very healthy cohort. This chemokine was also associated with a decrease in SIRT3 expression, a gene known to be important in aging and endothelial cell function.

To construct a chronic inflammation score, we used a guided autoencoder (GAE) algorithm, a deep learning method that efficiently deals with the network structure and non-linear behavior of the inflammatory response (Paul, 2013). Since deep learning algorithms can extract high-level complex abstractions as ‘data representations’ using non-linear functions, they are well-suited for the analysis of complex systems where most behaviors are non-linear, context dependent and organized in a distributed hierarchical fashion (Subramanian et al., 2015). In our case, this method outperformed other commonly used linear-modeling methods such as the elastic net (Zou, 2005), and PCA, and also a non-linear one such as plain autoencoder (Bengio, 2009) (Fig. S3b). The correlation between age and iAge was 0.78 (*P*<10^-16^), which is lower than that of the ‘proteomic age’ metric (R = 0.92), recently reported by Tanaka, et al. (Aging Cell, 2018). However, in contrast with the Tanaka study, which did not report disease associations, we find that iAge tracks with immunosenescence and multi-morbidity. In particular, we find a strong association with poor acute immune responses to cytokine stimuli, which is consistent with reports by two independent studies showing that high levels of baseline inflammatory markers correlate with weaker responses to hepatitis B and herpes zoster vaccine formulations (Fourati et al., 2016; Thevaranjan et al., 2017); and with our previous longitudinal studies of aging also showing that chronic inflammation is, at least in part, responsible for a reduced JAK-STAT response to cytokine stimulations in various leukocyte populations (Shen-Orr et al., 2016). Despite the proven utility of the cytokine stimulation assays used in our study with respect to an individual’s overall immune competence versus immune senescence (Furman et al., 2017a; Furman et al., 2013; Furman et al., 2015; Shen-Orr et al., 2016), a limitation of the assay relates to the stimuli used here which may not completely mirror the physiological stimuli that act on specific immune cell subsets *in vivo.* For example, while the stimuli we used strongly activate the memory compartment of bulk CD8 (+) and CD4 (+) T cells, these act relatively weakly on naïve T cells. Additional cell subsets that are poorly activated by the cytokines used in our study are Th1 cells, which can be activated by IL-12 and IL-18, or Th17 cells, which respond to other cytokine stimulations such as IL-1β or IL-18 in concert with IL-23 to produce Th-17 associated cytokines.

Inflammatory age was also positively correlated with multi-morbidity after controlling for age, high cholesterol, gender, BMI and CMV positivity. This suggests that this immune ‘metric’ for human health versus disease may be useful as a companion diagnostic to inform physicians about patient’s inflammatory status, especially those with chronic diseases. Furthermore, we find that in very healthy older adults with no clinical or laboratory evidence of cardiovascular disease, the largest contributor to inflammatory age, CXCL9, was positively correlated with subclinical levels of arterial stiffness and cardiac remodeling. These results strongly suggest that inflammatory age can also be used as an early molecular marker for cardiovascular malfunctioning.

CXCL9 is a T-cell chemoattractant induced by IFN-γ and mostly produced by neutrophils, macrophages and endothelial cells. Despite prior data showing that CXCL9 and other CXCR3 ligands are significantly elevated in hypertension and in patients with left ventricular dysfunction (Altara et al., 2015), here we find that in nominally healthy individuals, CXCL9 predicts subclinical levels of cardiovascular aging. At least two sources of CXCL9-mediated inflammation can ensue with aging based on our findings; one that is age-intrinsic and observed in aging endothelia, and one independent of age (likely as a response to cumulative exposure to environmental insults) and found in the validation cohort (Table S4). Interestingly, we did not find any significant correlation between known disease risk factors reported in the study (BMI, smoking, dyslipidemia), the levels of CXCL9 gene or protein expression. We thus hypothesize that one root cause of CXCL9 overproduction is cellular aging per-se, which triggers metabolic dysfunction - as was shown in many previous studies of aging - with production of damage-associated molecular patterns (DAMPs) such as adenine and N4-acetylcytidine (as demonstrated in our previous studies of aging, Furman D, 2017 Nat Med). These DAMPs can then act through the inflammasome machinery, such as NLRC4, to regulate multiple inflammatory signals including IL-1β (Furman, 2017) and CXCL9 (Janowski AM (2016) JCI).

Since endothelial cells but not cardiomyocytes expressed the CXCL9 receptor, CXCR3 (Fig. 3f), we hypothesize that this chemokine acts both in a paracrine fashion (when it is produced by macrophages to attract T cells to the site of injury) and in an autocrine fashion (when it is produced by the endothelium) to down-regulate SIRT3, an important stress-responsive deacetylase with cardio-protective and longevity enhancing properties involved in mitochondrial homeostasis, stem cells and tissue maintenance (Bonkowski and Sinclair, 2016; Lu et al., 2016). Of the 7 SIRT proteins, SIRT3 is the only member whose increased expression has been linked to the longevity of humans (Bellizzi et al., 2005; Rose et al., 2003), but the role of this protein in cardiovascular aging was not clear until recent observations in mice lacking the SIRT3 gene, which develop spontaneous cardiac hypertrophy and fibrosis at a young age (8 weeks-old) (Guo et al., 2017). Interestingly, the effects of SIRT3 in the heart are dependent on mitochondrial metabolism as shown in studies of SIRT3 KO mice, which have altered mitochondrial fatty acid oxidation (FAO) (Alrob et al., 2014; Hirschey et al., 2010) and decreased oxidative phosphorylation complex activity and ATP production (Ahn et al., 2008). Together this evidence and our results indicate that CXCL9 and SIRT3 play an important role linking inflammation, cell metabolism, endothelial cell function and cardiovascular remodeling, which is consistent with prior work showing intricate interactions between inflammation and cell metabolism in tissue repair processes (Eming et al., 2017).

Our data also place the endothelium as a central player in cardiovascular aging in agreement with Versari et al. (2009) who showed that cardiac hypertrophy is associated with endothelial dysfunction (Versari et al., 2009). They also suggest that endothelial cells may be one source of inflammation, but it is also possible that cardiomyocytes play a role since there is activation of the inflammasome NLRP3 in these cells in a model of acute myocardial infarction (Mezzaroma et al., 2011). Last, the hypertrophic response may also be affected by the adaptive immune response as was suggested both in non-ischemic heart failure (HF) patients and in mice with HF induced by transverse aortic constriction (TAC), where T cells exhibited enhanced adhesion to activated vascular endothelium. Moreover, T cell-deficient mice (TCRα(-/-)) treated with the TAC method had preserved left ventricular (LV) systolic and diastolic function, reduced LV fibrosis, hypertrophy and inflammation, and improved survival compared with wild-type mice (Nevers et al., 2015).

In summary, by analyzing immunological data from over 1000 individuals, we find a robust marker for chronic inflammation that could be potentially used to identify individuals with a decline in immunological function (immunosenescence), as well as those at risk for chronic disease and premature cardiovascular aging.

## Methods

### Study cohorts

#### The Stanford 1000 Immunomes Study Cohort

The Stanford 1KIP consists in 1001 ambulatory individuals (339 males and 662 females) recruited at Stanford University during the years 2007 to 2016 for various studies of aging and vaccination (*N* = 605) (Blazkova et al., 2017; Brodin et al., 2015; Furman et al., 2017b; Furman et al., 2014; Furman et al., 2013; Furman et al., 2015; Price et al., 2013; Roskin et al., 2015; Shen-Orr et al., 2016; Wang et al., 2014), and for an independent study of chronic fatigue syndrome (Montoya et al., 2017), from which we utilized data from the control set of participants only (*N* = 396).

#### Aging and vaccination study cohort

Study participants were enrolled in an influenza vaccine study at the Stanford-LPCH Vaccine Program during the years 2007 to 2016. Since baseline samples were obtained from all the individuals prior to vaccination with the influenza vaccine, no randomization or blinding was done for this study. The protocol for this study was approved by the Institutional Review Board of the Research Compliance Office at Stanford University. Informed consent was obtained from all subjects. All individuals were ambulatory. At the time of initial enrollment volunteers had no acute systemic or serious concurrent illness, no history of immunodeficiency, nor any known or suspected impairment of immunologic function, including clinically observed liver disease, diabetes mellitus treated with insulin, moderate to severe renal disease, blood pressure >150/95 at screening, chronic hepatitis B or C, recent or current use of immunosuppressive medication. In addition, on each annual vaccination day, none of the volunteers had been recipients or donors of blood or blood products within the past 6 months and 6 weeks respectively, and none showed any signs of febrile illness on day of baseline blood draw. Peripheral blood samples were obtained from venipuncture and mononuclear cells were separated and stored at the Stanford Clinical & Translational Research Unit (CTRU). Whole blood was used for gene expression analysis (below). Serum was separated by centrifugation of clotted blood, and stored at ^-^80°C before CMV serology, cytokine and chemokine determination.

#### Chronic Fatigue Syndrome Study cohort

Study participants were recruited from Northern California from March 2, 2010 to September 1, 2011. Their peripheral blood was drawn between 8:30 AM and 3:30 PM on the day of enrollment. These samples were collected at baseline for each participant (no exercise prior to blood sampling). In addition, as each ME/CFS patient was being recruited into the study, two corresponding, age and sex matched controls, were contemporaneously enrolled until the target sample size of 200 patients and 400 controls was achieved. This approach resulted in patients and controls being intercalated in their time of entry into the study. Eight milliliters of blood were drawn into a red-topped serum tube (Fisher Scientific) by the CTRU’s phlebotomy team. Serum was obtained by allowing blood to clot for 40 min. Once clotted, the blood tube was centrifuged in a refrigerated (4°C) centrifuge (Allegra X-15R, Beckman Coulter) at 2000xG for 15 minutes. Serum was isolated and mixed thoroughly in a tube using a 2-milliliter sterile, serological pipette (Fisher Scientific) to obtain a homogenous solution prior to dispensing to storage tubes. Serum was distributed into aliquots per the Stanford Human Immune Monitoring Center (HIMC, http://iti.stanford.edu/himc.html) aliquot guidelines and frozen at −80°C. For the day of the cytokine assay, matched sets of ME/CFS cases and healthy controls were always mixed in all plates to reduce confounding case status with plate artifacts. In summary, ME/CFS patients and controls were treated identically in terms of recruitment and sera handling protocols.

#### Validation cohort and centenarians

A total of 37 subjects were enrolled by two Italian study centers (Bologna and Florence) and surrounding areas. The group of centenarians consisted of 19 subjects (10 men, mean age 102.8 ± 2.3 years and 9 women, mean age 103.7 ± 2.6 years) and the group of controls consisted of 18 subjects (9 men, mean age 64.8 ± 7.9 years and 9 women, mean age 67.1 ± 7.3 years). The lists of subjects were obtained by the Office of Vital Statistics. All participants signed the informed consent before undergoing the questionnaires (functional and cognitive status, depression, self-perceived health), measurements (anthropometric measures, blood pressure, physical performance) and blood sampling. History of past and current diseases was accurately collected by checking the participants’ medical documentation and addressing the major age-related pathologies. Current use of medication (including inspection of the drugs by the interviewer) was recorded. The study protocol was approved by the Ethical Committee of Sant’Orsola-Malpighi University Hospital (Bologna, Italy). Overnight fasting blood samples were obtained in the morning. Plasma was obtained within 2 hours from venipuncture by centrifugation at 2000 g for 20 min at 4°C, rapidly frozen and stored at −80°C.

#### Cardiovascular study cohort

After approval by Stanford’s Institutional Review Board, 151 subjects participating in the National Institute of Health sponsored 5 U19 AI05086 study and Stanford Cardiovascular Institute Aging Study were screened for inclusion in the study. The screening process included a comprehensive health questionnaire including the London School of Hygiene Cardiovascular questionnaire. Exclusion criteria included the following: history of acute or chronic illness such as atherosclerosis, systemic hypertension, diabetes mellitus or dementia, familial history of early cardiovascular disease (<55 year old), on non-steroidal anti-inflammatory drugs or on inhaled steroids on a regular basis, history of malignancies, history of surgery within the last year, history of atopic skin disease, history of infection within the last 3 months including upper respiratory infections or urinary infections, and history of vaccination within the past 3 months. Patients older than 80 years old who had a prior history of mild systemic hypertension but with a normal blood pressure at the time of the visit (blood pressure < 140/90 mmHg) were not excluded from the study. Based on the inclusion and exclusion criteria, 97 subjects were included in the study. We divided the patients into 4 groups according to pre-specified age boundaries (i.e., between 25 and 44, 45 to 60, 60 to 75 and 75 to 90 years old).

### Experimental Procedures

#### Human iPSC Generation and Culture

Protocols for isolation and use of patient blood were approved by the Stanford University Human Subjects Research Institutional Review Board. Details on the isolation of patient blood, as well as the generation, culture, characterization, of hiPSCs, have been previously published (Matsa et al., 2016).

#### Human iPSC Differentiation to Endothelial Cells

Human iPSCs (hiPSCs) were seeded on matrigel plates and grown in hiPSC medium for 4 days to 75-80% confluency. Differentiation to ECs were initiated by treating the hiPSCs with 6 μM CHIR99021 in RPMI-B27 without insulin media (Life Technologies) for 2 days, followed by another treatment of 2 μM CHIR99021 in RPMI-B27 without insulin media for 2 days. Following these treatments, the differentiating hiPSCs were subjected to endothelial medium EGM2 (Lonza) supplemented with 50 ng/mL VEGF, 20 ng/ml BMP4, and 20 ng/ml FGFb for 7 days, with medium being changed every 2 days. On day 12, induced ECs were isolated using MAC sorting, where cells were first dispersed by trypsin, then incubated with CD144 antibody, and finally passed through a MACS column containing CD144-conjugated magnetic microbeads (Miltenyi Biotec). The sorted cells were then seeded on 0.2% gelatin-coated plates and maintained in EGM2 media supplemented with 10 µM SB431542 (TGFb inhibitor). hiPSC-ECs were passaged on confluence and maintained in EGM2 medium.

#### *In vitro* monolayer cardiomyocyte differentiation of human iPCSs

To induce cardiomyocyte differentiation, approximately 1×10^5^ undifferentiated hiPCSs were seeded in each well of Matrigel-coated 6-well plates and cultured in differentiation medium following previous protocol (Gu et al., 2017; Lian et al., 2012). Glucose-free MEMα supplemented with FBS and lactate has been employed to enrich cells to 98.0% α-actinin–positive at 37°C, 20% O2, and 5% CO_2_ in a humidified incubator with medium changes every 48 hours, and cells were passaged once they reached 80-90% confluence. Human iPSC-CMs were treated immediately after enrichment.

#### Cell lines

Human Umbilical Vein Endothelial Cells (HUVEC) were purchased from Lonza and cultured following their protocol. Fibroblasts were cultured as previously described (Takahashi and Yamanaka, 2006).

#### Cardiovascular phenotyping

Cardiovascular age was assessed using 3 parameters: (1) aortic pulse wave velocity (PWV), a measure of vascular stiffness; (2) relative wall thickness (RWT), a measure of ventricular remodeling, and (3) early diastolic mitral annular velocities (e’), a measure of ventricular relaxation. In addition, we measured the ratio of early mitral inflow velocity (E) to e’, a surrogate marker of end-diastolic filling pressures (Nagueh et al., 2009; Redfield et al., 2005).

Aortic PWV was calculated as the ratio of the pulse wave distance (in meters) to the transit time (in seconds). A 9.0 MHz Philips linear array probe was used to assess the carotid arteries (main common, bulb and internal carotid artery) and proximal femoral arteries. Pulse wave distance (D) was measured as the distance from the sternal notch to the femoral artery (x_direct_) from which we subtracted the distance from the sternal notch to proximal descending aorta (D = x_direct_ - x_notch-aota_). The intersecting tangent method was used to measure the time from a reference ECG signal and the foot of the pulse wave. Heart rate had to be within 2 bpm between the carotid and femoral signal. All Doppler signals were recorded at 150 mm/s. Inter-observer variability was calculated on 50 samples in our laboratory and the intraclass correlation coefficient was 0.94 for PWV measured by 2 independent observers (independent measures of path length and transit time).

Echocardiograms were performed using the Philips IE33 echocardiographic system according to the recommendations of the American Society of Echocardiogram.(Lang et al., 2005) All studies were interpreted by one physician (FH) who was blinded to age as well as clinical and biological data. All parameters were measured in triplicate and averaged. Ventricular dimensions and wall thickness were measured using M-mode derived measures; we excluded septal band from the measurement of the septum and chordates from the measurements of the posterior wall. RWT was calculated as the sum of septal and posterior wall divided by left ventricular internal dimensions. Ventricular mass was estimated using the ASE recommended formula based on modeling the LV as a prolate ellipse (Lang et al., 2005). Left ventricular ejection fraction was estimated using the Simpson biplane method. The tissue Doppler e’ velocity represents an average of the septal and lateral annulus (Nagueh et al., 2009; Redfield et al., 2005). Inter-observer variability was calculated on 50 samples; the intraclass correlation in our laboratory is 0.93 for LV mass measurements.

#### Induced human pluripotent stem cell-derived cardiomyocytes and endothelial cells

We derived hiPCSs from 5 healthy individuals and cell lines passed common assessments for pluripotency and genomic stability. These hiPCSs were differentiated into cardiomyocytes to purities of >85% and endothelial cells to purities of > 90%. hiPCS-CMs were differentiated on day 30 and hiPCS-ECs were differentiated on day 14. Both types of cells expressed mature cell markers. The experimental component of our study focused on 3 different components of INF-γ signaling pathway.

To analyze gene expression CXCR3, RNA was isolated using an RNeasy Plus kit (QIAGEN), cDNA was produced using a High Capacity RNA-to-cDNA kit (Life Technologies), and real-time PCR was performed using TaqMan Gene Expression Assays, TaqMan Gene Expression Master Mix, and a StepOnePlus^TM^ Real-Time PCR System (Life Technologies). All PCR reactions were performed in triplicate, normalized to the GAPDH endogenous control gene, and assessed using the comparative Ct method.

#### Quantitative real-time PCR

To analyze the gene expression pattern for CXCL9 and SIRT3, RNA was extracted using Qiagen RNA isolation kit (Qiagen 74104) and cDNA was synthesized using qScript cDNA SuperMix (QuantaBio). Real-time PCR was performed using TaqMan Gene Expression Assays (GAPDH, Hs02758991_g1, CXCL9, Hs00171065_m1, SIRT3, Hs00953477_m1), TaqMan Master Mix, and a 7900HT Real-Time PCR System (ThermoFisher Scientific). All PCR reactions were performed in triplicate, normalized to the GAPDH housekeeping gene, and assessed using the ΔΔCt relative quantification (RQ) method.

#### Vascular Tube-like Formation

The functions of the generated hiPSC-ECs were characterized in angiogenic assays and compared to hiPSCs. The generated hiPSC-EC were assessed for their ability to form tubelike structures by seeding 1 x 10^4^ cells in wells coated with Matrigel (Corning® Matrigel® Matrix) containing EGM2 medium supplemented with 50 ng/mL VEGF and incubated for 16-24 hours.

#### Isometric Tension Recordings

Mouse thoracic aortas were carefully dissected, and the vessels were transferred to a dish with ice-cold Krebs Solution (in mmol/L: 133 NaCl, 4.6 KCl, 2.5 CaCl2,16.3 NaHCO3, 1.75 Na2HPO4, 0.6 MgSO4, 10 glucose). The vessels were cut into small rings and mounted on an isometric wire myograph chambers (Danish Myo Technology) and subjected to a normalization protocol. Following normalization, the vessels were incubated with either PBS or different concentrations of recombinant mouse CXCL9 protein (R&D systems, catalog number 492-MM) for 3-4 hrs. A concentration-dependent contraction curve was created by accumulative application of the prostaglandin agonist U46619. Subsequently, concentration-dependent relaxation curves of Acetylcholine were conducted on these vessels and percent relaxation calculated for each dose.

#### CXCL9 knockdown

Gene knockdown experiments were performed using GIPZ CXCL9 shRNA Viral Particle Starter Kit (Dharmacon Inc, CO) containing a pool of select shRNA. These included V3LHS_368350 (TAGACATGTTTGAACTCCA), V3LHS_409682 (AGTTATATACTGTCTACCT), V3LHS_409683 (AGAAGAACAAAGACAATCA). The MOI (multiplicity of infection) of CXCL9 shRNA was assessed after 72 hours infection by puromycin selection and GFP analysis according to manufacturer’s instructions. iPSCs were transfected with CXCL9 knockdown lentivirus at MOI >0.9, and knockdown efficiency was measured by real-time quantitative RT-PCR.

#### Flow Cytometry Immunophenotyping

This assay was performed by the Human Immune Monitoring Center at Stanford University. PBMC were thawed in warm media, washed twice and resuspended at 1×10^7^ viable cells/mL. 50 uL cells per well were stained for 45 min at room temperature with the antibodies shown in the Key Resources Table (all reagents from BD Biosciences, San Jose, CA). Cells were washed three times with FACS buffer (PBS supplemented with 2% FBS and 0.1% sodium azide), and resuspended in 200 uL FACS buffer. 100,000 lymphocytes per sample were collected using DIVA 6.0 software on an LSRII flow cytometer (BD Biosciences). Data analysis was performed using FlowJo v9.3 by gating on live cells based on forward versus side scatter profiles, then on singlets using forward scatter area versus height, followed by cell subset-specific gating.

#### Phosphoepitope Flow Cytometry (Cytokine stimulation, pSTAT readouts)

This assay was performed by the Human Immune Monitoring Center at Stanford University. PBMC were thawed in warm media, washed twice and resuspended at 0.5×10^6^ viable cells/mL. 200 uL of cells were plated per well in 96-well deep-well plates. After resting for 1 hour at 37°C, cells were stimulated by adding 50 ul of cytokine (IFNa, IFNg, IL-6, IL-7, IL-10, IL-2, or IL-21) and incubated at 37°C for 15 minutes. The PBMCs were then fixed with paraformaldeyde, permeableized with methanol, and kept at −80C overnight. Each well was bar-coded using a combination of Pacific Orange and Alexa-750 dyes (Invitrogen, Carlsbad, CA) and pooled in tubes. The cells were washed with FACS buffer (PBS supplemented with 2% FBS and 0.1% soium azide), and stained with the following antibodies (all from BD Biosciences, San Jose, CA): CD3 Pacific Blue, CD4 PerCP-Cy5.5, CD20 PerCp-Cy5.5, CD33 PE-Cy7, CD45RA Qdot 605, pSTAT-1 AlexaFluor488, pSTAT-3 AlexaFluor647, pSTAT-5 PE. The samples were then washed and resuspended in FACS buffer. 100,000 cells per stimulation condition were collected using DIVA 6.0 software on an LSRII flow cytometer (BD Biosciences). Data analysis was performed using FlowJo v9.3 by gating on live cells based on forward versus side scatter profiles, then on singlets using forward scatter area versus height, followed by cell subset-specific gating.

#### CyTOF Immunophenotyping

This assay was performed in the Human Immune Monitoring Center at Stanford University. PBMCs were thawed in warm media, washed twice, resuspended in CyFACS buffer (PBS supplemented with 2% BSA, 2 mM EDTA, and 0.1% soium azide), and viable cells were counted by Vicell. Cells were added to a V-bottom microtiter plate at 1.5 million viable cells/well and washed once by pelleting and resuspension in fresh CyFACS buffer. The cells were stained for 60 min on ice with 50 uL of the following antibody-polymer conjugate cocktail depicted in the Key Resources Table. All antibodies were from purified unconjugated, carrier-protein-free stocks from BD Biosciences, Biolegend, or R&D Systems. The polymer and metal isotopes were from DVS Sciences. The cells were washed twice by pelleting and resuspension with 250 uL FACS buffer. The cells were resuspended in 100 uL PBS buffer containing 2 ug/mL Live-Dead (DOTA-maleimide (Macrocyclics) containing natural-abundance indium). The cells were washed twice by pelleting and resuspension with 250 uL PBS. The cells were resuspended in 100 uL 2% PFA in PBS and placed at 4C overnight. The next day, the cells were pelleted and washed by resuspension in fresh PBS. The cells were resuspended in 100 uL eBiosciences permeabilization buffer (1x in PBS) and placed on ice for 45 min before washing twice with 250 uL PBS. If intracellular staining was performed, the cells were resuspended in 50 uL antibody cocktail in CyFACS for 1 hour on ice before washing twice in CyFACS. The cells were resuspended in 100 uL iridium-containing DNA intercalator (1:2000 dilution in PBS; DVS Sciences) and incubated at room temperature for 20 min. The cells were washed twice in 250 uL MilliQ water. The cells were diluted in a total volume of 700 uL in MilliQ water before injection into the CyTOF (DVS Sciences). Data analysis was performed using FlowJo v9.3 (CyTOF settings) by gating on intact cells based on the iridium isotopes from the intercalator, then on singlets by Ir191 vs cell length, then on live cells (Indium-LiveDead minus population), followed by cell subset-specific gating.

#### Phosphoepitope CyTOF (Cytokine stimulation, pSTAT readouts)

This assay was performed by the Human Immune Monitoring Center at Stanford University. PBMC were thawed in warm media, washed twice, counted by Vi-cell and resuspended at 5×10^6^ viable cells/mL. 100 uL of cells were plated per well in 96-well deep-well plates. After resting for 1 hour at 37°C, cells were stimulated by adding 25 ul of stim (IFNa, IL-6, IL-7, IL-10, IL-21, LPS or PMA/ionomycin) and incubated at 37°C for 15 minutes. Cells were then fixed with paraformaldehyde, washed twice with CyFACS buffer (PBS supplemented with 2% BSA, 2 mM EDTA, and 0.1% sodium azide) and stained for 30 min at room temperature with 20 uL of surface antibody cocktail. Cells were washed twice with cyFACS, permeabilized with 100% methanol and kept at −80C overnight. Next day cells were washed with cyFACS buffer and resuspended in 20 uL intracellular antibody cocktail in CyFACS for 30 min at room temperature before washing twice in CyFACS. Cells were resuspended in 100 uL iridium-containing DNA intercalator (1:2000 dilution in 2% PFA in PBS) and incubated at room temperature for 20 min. Cells were washed once with cyFACS buffer and twice with MilliQ water. Cells were diluted to 750×10^5 cells/mL in MilliQ water before injection into the CyTOF. Data analysis was performed using FlowJo v9.3 (CyTOF settings) by gating on intact cells based on the iridium isotopes from the intercalator, then on singlets by Ir191 vs cell length followed by cell subset-specific gating.

#### Determination of Serum Immune Proteins

Luminex -Polystyrene bead kits. This assay was performed in the Human Immune Monitoring Center at Stanford University. Human 50 or 51-plex kits were purchased from Panomics/Affymetrix and used according to the manufacturer’s recommendations with modifications as described below. Briefly, samples were mixed with antibody-linked polystyrene beads on 96-well filter-bottom plates and incubated at room temperature for 2 h followed by overnight incubation at 4°C. Room temperature incubation steps were performed on an orbital shaker at 500-600 rpm. Plates were vacuum filtered and washed twice with wash buffer, then incubated with biotinylated detection antibody for 2 h at room temperature. Samples were then filtered and washed twice as above and re-suspended in streptavidin-PE. After incubation for 40 minutes at room temperature, two additional vacuum washes were performed, and the samples re-suspended in Reading Buffer. Each sample was measured in duplicate. Plates were read using a Luminex 200 instrument with a lower bound of 100 beads per sample per cytokine. Custom assay Control beads by Radix Biosolutions are added to all wells.

#### SOMAscan Assay

Thirty-seven plasma samples (19 males and 18 females) from 2 different cohorts (PRIN06 and PRIN09) were used in this study. Samples were stored at −80 °C and sent on dry ice to SomaLogic Inc. (Boulder, Colorado, US). The SOMAscan platform was used to quantify levels of plasma proteins. Assay details have been previously described (Gold et al., 2010). Briefly, this platform is based on modified single-stranded DNA (SOMAmers) that are used to bind to specific protein targets. Data in Relative Fluorescent Units (RFU) for 1305 SOMAmer probes were obtained for these samples and no samples or probe data was excluded. PRIN06 and PRIN09 samples were measured in 2 batches. Datasets were bridged to each other using Somalogic calibrators.

### Quantification and Statistical Analysis

#### Normalization procedures for Luminex assays

We conducted a 3-step normalization procedure. First, we used an internal control (CON-S) that was run on each batch to plate-normalize the data. We considered the regression model

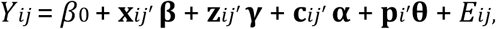

where outcome is the cytokine’s median fluorescence intensity averaged over duplicate wells (aMFI) for *j*th participant on *i*th plate, **x** is the design vector of variables of interest with corresponding regression coefficients **β**, and **z** is the design vector of nuisance variables of corresponding regression coefficients **γ**. These may include baseline covariates, random coefficients to model longitudinal data, etc. Covariance among repeated observations within participants (e.g., longitudinal aMFI) was modeled via *Cov*[*Eijk, Eijk′*] for *k* ≠ *k′*. Parameter *β*0 is the intercept. This model also adjusts for nonspecific binding **c** (used CHEX4) (with corresponding regression coefficients **α**) (Montoya et al., 2017). Vector form permits modeling of any nonlinear effects of nonspecific binding on outcome. For plate effects, we used the indicator-variable vector **p** of plate effects has regression coefficients **θ**. The variance of regression residual *E* is allowed to vary among plates, such as Var[*Eij*] ≠ Var[*Ei′j*] for *i* ≠ *i*′. Together, **p**′**θ** and variance Var[*Eij*] account respectively for location and scale effects of plates. Secondly, for source normalization (Influenza vaccine studies versus chronic fatigue study), we used a naïve correction in which principal component analysis is conducted on all the data and the effect of top components is removed by regression analysis on data source until the batch effect is no longer significant. The mean absolute correlation is then computed as a function of the number of PCA components in batch corrected vs. raw data and heat-maps for before and after PCA-correction are also shown (Fig. S5).

#### Guided Auto-Encoder (GAE) and inflammatory age

When dealing with data with a large number of dimensions and complex network structure, we aimed to find a non-linear method to summarize the data possibly to a compact representation. This compact representation can be further used for feature extraction, visualization, or classification purpose. To obtain the informative representation, we proposed a novel model called “Guided Auto-Encoder”. The method is built based on Auto-Encoder with a combined objective. Auto-encoders use a non-linear transformation of the data and hence, it can model complex processes (Bengio, 2009). One problem of auto-encoders is re-parameterization. With different initialization, it could have different results. Among the different types of visualizations with similar summarization level, one usually wants a representation that is informative of a specific target. Hence, we can construct a representation with two focuses: 1) the learned compact representation h can be recovered to the original data as much as possible (reconstruction loss); 2) the learned compact representation should be as informative of the desired target as possible (prediction loss). Therefore, we proposed a novel structure – guided-auto-encoder –that balances the two objectives in order to provide an informative representation. We applied the GAE to extract an immunology score or inflammatory age. It is a non-linear transformation of the cytokine data in a person that both approximates the true age, while preserves the information of the cytokine level.

#### Auto-encoder

Given the input data vector x, an auto-encoder aims to reconstruct the input data vector x. We consider an auto-encoder with L encoding layers and L decoding layers has depth of L, and each layer has fixed number of hidden nodes m.

For convenience, the input layer is defined as h_0_(x) = x, and the output of lth hidden layer is defined as h_l_(x). The number of nodes in layer l is m_l_. The input into the lth layer of the network is defined as:

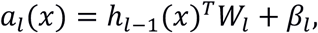

where W_l_ is a real value weight matrix of m_l-1_ by m_l_ and β_l_ is a vector of length m_l-1_. The output of lth hidden layer is:

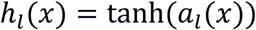

where tanh is the hyperbolic tangent function:

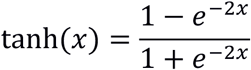

We define the output of the Lth layer h_L_(x) as the coding layer. The decoding layers are from L+1 to 2L-1 layer with the same setting. Finally, a linear output layer is on top of the last decoding layer:

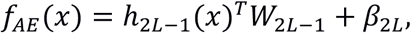

Given data vectors x, we train an auto-encoder, we minimize the reconstruction loss on the data:

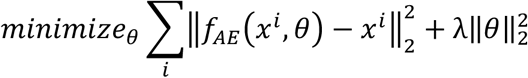

where i ranges of the number of samples, *θ* represents all the parameters used in the auto-encoder, and λ is the weight decay penalty used for regularization. To optimize the objective (1), we used a stochastic optimization method ADAM (Kingma and Jimmy, 2014).

### Guided-Auto-encoder

A guided auto-encoder aims to reduce both the reconstruction loss and predictive loss. Given the input x, a side-phenotype y and an auto-encoder *f_AE_*, the guided-auto-encoder incorporates a predictive function on the coding layer:

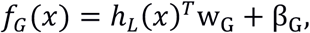

with its own set of parameters w_G_ and β_G_

Let *θ* be the set of all parameters of a GAE, the training objective is:

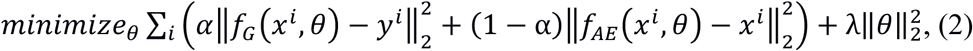

where is **α** real value number between 0 and 1 that is called the *guidance-ratio*. An example guided-auto-encoder with depth 2 and width 3 is shown below.

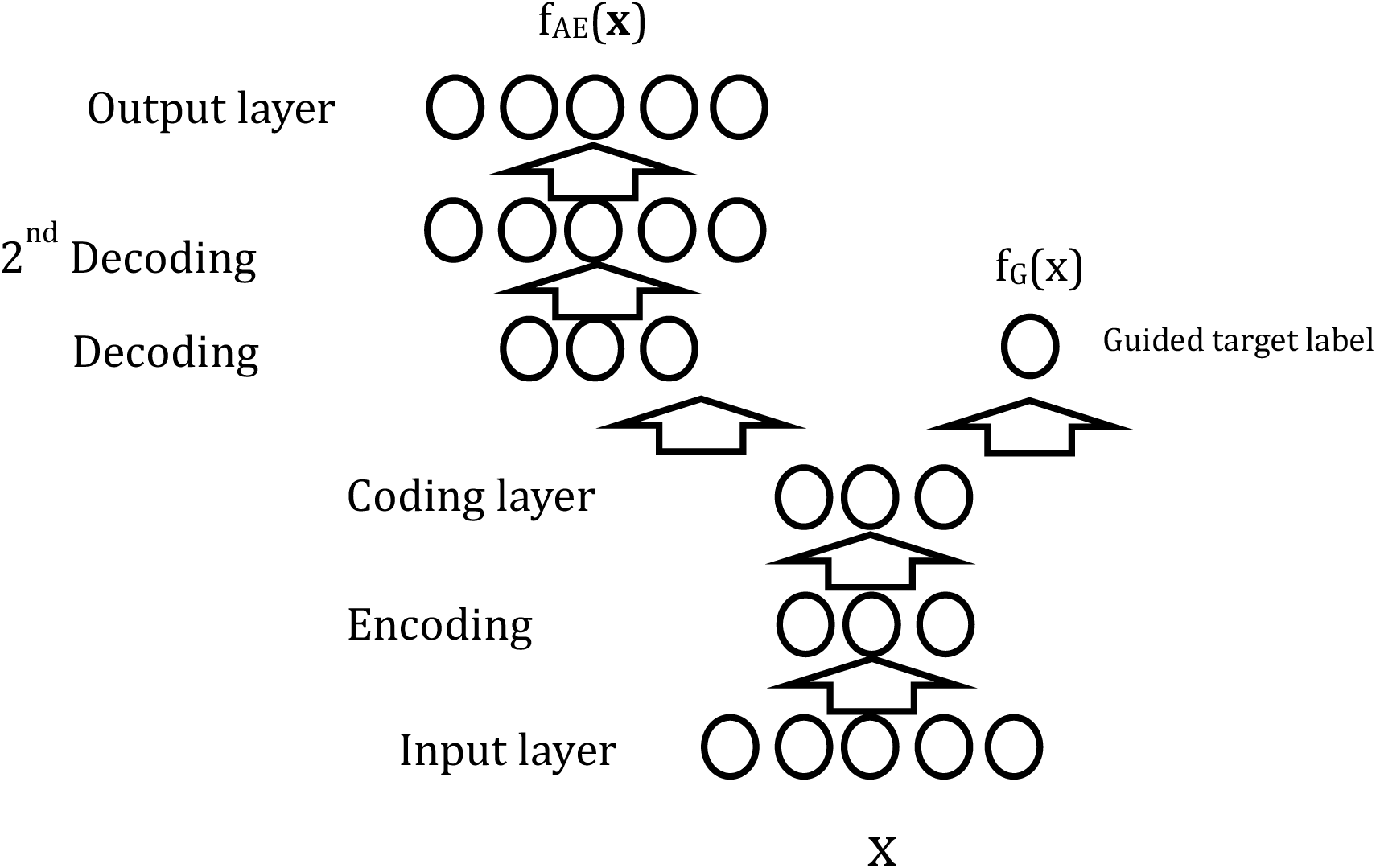

We use optimization method ADAM (Kingma and Jimmy, 2014) to minimize objective. By choosing different guidance-ratio, we can reach different level of balance between prediction loss and reconstruction loss.

#### Extraction of Inflammatory age

In order to provide a marker summarization of a patient’s immune system health state, we invented a novel quantity – inflammatory age. This quantity is age of patient predictable from the inflammatory state of the immune system. In order to obtain this quantity we focused on cytokine measurements. By construction, the inflammatory age is a non-linear function of cytokine measurements, but also an estimate of the patient’s true age.

To construct this quantity, we used Guided Auto-Encoder (GAE) aimed to compactly represent cytokine measurements and predict side-phenotype chronological age. We identified best code length, among lengths from 1 to 10, using a 5-fold-cross-validation. We select the length of code k, whose performance was not statistically significantly worse than that of longer codes (paired t-test p-value>0.05). Within each fold we performed nested 3-fold cross-validation to select hyper-parameters (depth, weight decay and guidance-ratio).

After obtaining the best code length as 5 (Fig. S3a), we used the 5-fold-cross-validation to select the best hyper-parameter setting (depth = 2, guidance-ratio = 0.2, L2 = 0.001) on all GAE with code length 5. Finally, we trained the GAE on the whole dataset with the selected best hyper-parameter setting and obtained the predictive function as the inflammatory age predictor. To derive the inflammatory index, we subtracted chronological age from inflammatory age.

#### Prediction of multi-morbidity using cyclical coordinate descent and correlation with immunosenescence

We hypothesized that important immune components will emerge from fitting a linear regression model with l1 and l2 penalties, the elastic net penalty, a regularization algorithm that uses cyclical coordinate descent in a pathwise fashion as described (Friedman et al, 2007, 2010). We envisioned an unbiased approach to select predictors of multi-morbidity based on available data for all 902 subjects while controlling for the age effect. A total of 127 features were included in the prediction model. We assume all of our predictors are standardized to mean 0 and s.d. 1. The result of our fitting procedure is the set of predictor weights β and intercept α for the linear regression model. In practice, penalty weights are set by a data-driven procedure, such as 10-fold cross validation. The minimum λ was chosen to yield the lowest MAE with the minimum set of features. We envisioned age-controlled feature selection by imposing feature-specific penalty. In this procedure feature age is ‘forced in’ the model and l1 penalty is let to choose from all other features. In practice, we create a vector of size 127 and chose α = 0 for feature age and α = 1 for the remainder 126 features. To investigate the effect of iAge on immunosenescence, multiple regression analysis was conducted using iAge as a predictor variable (controlled by age, sex, CMV) and the frequency of naïve CD8 (+) T cells, as a target variable (surrogate of immunosenescence). Similarly, the effect of iAge (after controlling for age, sex and CMV) was estimated on a total of 92 cell stimulations. Adjusted p-values (by permutation tests) were combined by using a modified Fisher’s combined probability test (Dai et al., 2014).

#### Estimation of Inflammatory Age in centenarians and older control cohorts

We first aimed at estimating the minimum set of features required for accurate prediction of inflammatory age. To do so, we used the results from our previous analysis in which we investigated the composition of inflammatory age based on the first order partial derivative of inflammatory age (jacobians). We sorted the immune features based on their absolute jacobian and subsequently generated *n* - feature set models, each with a different feature number and removed one feature at a time starting from the less to the most important, which is never fully discarded. Since one important immune protein was not measured in the SOMAscan assay (the TNF-related apoptosis-inducing ligand (TRAIL)), we aimed at building a regression model to predict inflammatory excluding TRAIL that yields the same accuracy as model 1. Hence, we compared the inflammatory age prediction accuracy of model 1, to the accuracy of a series of models (TRAIL excluded) including an increasing number of features based on feature contribution to inflammatory age, as done before. Using the Stanford 1KIP data, we found that the prediction accuracy of a model when TRAIL is removed but containing EOTAXIN, MIP-1α, CXCL1, CXCL9, IFN-γ, IL-1β, IL-2, LEPTIN and PAI-1 was not different from that of the model 1 (in which TRAIL is included) (by likelihood ratio test, *P* < 0.01). We then directly estimated inflammatory age as the predicted age of subjects in the aging control cohort and centenarians based on standardized coefficients from the previous analysis on the Stanford 1KIP dataset and normalized RTU values for EOTAXIN, MIP-1α, CXCL1, CXCL9, IFN-γ, IL-1β, IL-2, LEPTIN and PAI-1. Inflammatory age was then used to compute inflammatory index (inflammatory age minus chronological age) in these cohorts.

## Data and Software Availability

### Data availability

The cell subpopulation, immune protein and cell signaling data for the Stanford Aging and Vaccination studies are publicly available on ImmPort Bioinformatics Repository (http://www.immport.org/immport-open/public/home/home) under the following study IDs SDY311 (cytokines, phosphoflow assays and CyTOF surface phenotyping), SDY312 (cytokines, phosphoflow assays and flow cytometry surface phenotyping), SDY314 (flow cytometry surface phenotyping), SDY315 (cytokines, phosphoflow assays and CyTOF surface phenotyping) and SDY478 (cytokines and CyTOF surface phenotyping).

### Software availability

Maximum likelihood estimation (MLE) is a function of the STATS4 R package that can be found under the following link: https://www.rdocumentation.org/packages/stats4/versions/3.4.1. The code used for the identification of immunotypes and construction of inflammatory age has been deposited on GitHub (https://github.com/) and is available under the following link: https://github.com/clingsz/GAE. For the immunological characterization of immunotypes we used R programming (https://www.r-project.org/). The Lasso and Elastic-Net regularized generalized linear models package (glmnet) for R programming can be found at the following link: https://cran.r-project.org/web/packages/glmnet/index.html

## Acknowledgments

We thank the study participants for their time and dedication, and the staff of the Stanford-LPCH Vaccine Program for recruiting participants and conducting the studies. We are also thankful to Professor Mark M. Davis for his invaluable ideas and insightful discussions.

Support for the conduct of these studies was from The Ellison Foundation, NIH U19 AI057229, U19 AI090019, and NIH/NCRR CTSA award number UL1 RR025744. CT.gov numbers for the vaccine studies are NCT01827462, NCT02133781, NCT03020498, NCT03020537, NCT01987349, NCT03022396, NCT03022422, NCT03022435, NCT03023176 and NCT02141581. This work was also supported by grants to C.F. from the European Union (EU) Horizon 2020 Project PROPAG-AGEING (grant 634821), the EU JPND ADAGE project, the Ministry of Education and Science of the Russian Federation Agreement (grant 074-02-2018-330). We gratefully acknowledge additional funding support from the National Institutes of Health (NIH) K01 HL135455, Stanford TRAM scholar award (N.S.), the Paul F. Glenn Foundation and the NIH Stanford Alzheimer’s Disease Research Center P50AG047366.

## Author Contributions

Conceptualization, D.F.; Methodology, D.F., F.H., R.T., T.H., H.T.M., and C.L.D.; Software, D.F., T.G., V.J., Validation, F.H., R.O., D.M., J.C.W., and C.F.; Formal Analysis, D.F., T.G., N.S., B.L., T.K., and V.J.; Investigation, D.F., L.C., N.S., Y.R-H., and B.L.; Resources, F.H., T.K., R.O., D.M., B.L., H.T.M., C.L.D., T.W-C., C.F., V.J., J.C.W., J.G.M.; Writing – Original Draft, D.F; Writing – Review & Editing, D.F., N.S., F.H., L.C., T.K., S. S-O., H.T.M., C.L.D., C.F., J.C.W., J.G.M.; Visualization, D.F., and T.G., Supervision, F.H., R.T., T.H., H.T.M., C.L.D., T.W-C., C.F., V.J., J.C.W., J.G.M.; Project administration, R.O., D.M., H.T.M., C.L.D., and J.G.M.; Funding Acquisition, D.F., C.F., J.C.W., and J.G.M.

## Conflict of interests

The authors declare no conflict of interests.

## Supplementary Figures

**Supplementary Figure 1.**
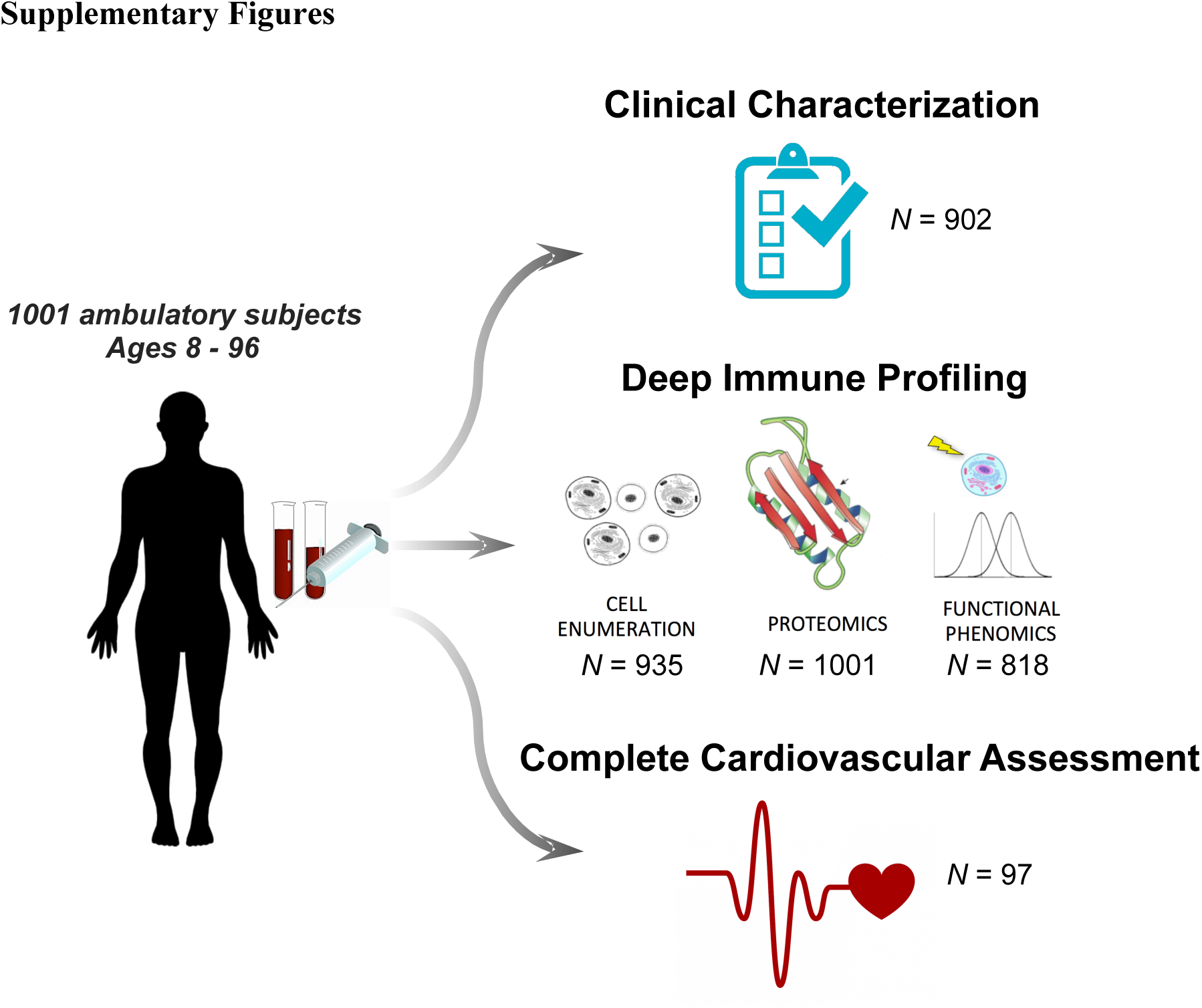
1000 Immunomes Study design: systematic analysis of immune systems via ‘OMICS’ approaches. The Stanford 1000 Immunomes Project consist of 1001 ambulatory subjects age 8 to >89 (34% males, 66% females) recruited during the years 2007 to 2016 for a longitudinal study of aging and vaccination (Blazkova et al., 2017; Brodin et al., 2015; Furman et al., 2017; Furman et al., 2014; Furman et al., 2013; Furman et al., 2015; Price et al., 2013; Roskin et al., 2015; Shen-Orr et al., 2016; Wang et al., 2014), and for an independent study of chronic fatigue syndrome (Montoya et al., 2017) from which only healthy controls were included. For all samples of the Stanford 1KIP, deep immune phenotyping was conducted at the Stanford Human Immune Monitoring Center, where peripheral blood specimens were isolated and analyzed using standard procedures. Peripheral blood samples were obtained by venipuncture and peripheral blood mononuclear cells or whole blood samples were used for determination of cellular phenotypes and frequencies (*N* = 935) and for investigation of *in vitro* cellular responses to a variety of cytokine stimulations (*N* = 818); serum samples were obtained and used for protein content determination (including a total of 50 cytokines, chemokines and growth factors) (*N* = 1001). Clinical characterization was assessed via clinical questionnaire in a total of 902 subjects who completed the full set of 53 clinical items. From a total of 97 healthy young and older adults, comprehensive cardiovascular phenotyping was also conducted.

**Supplementary Figure 2.**
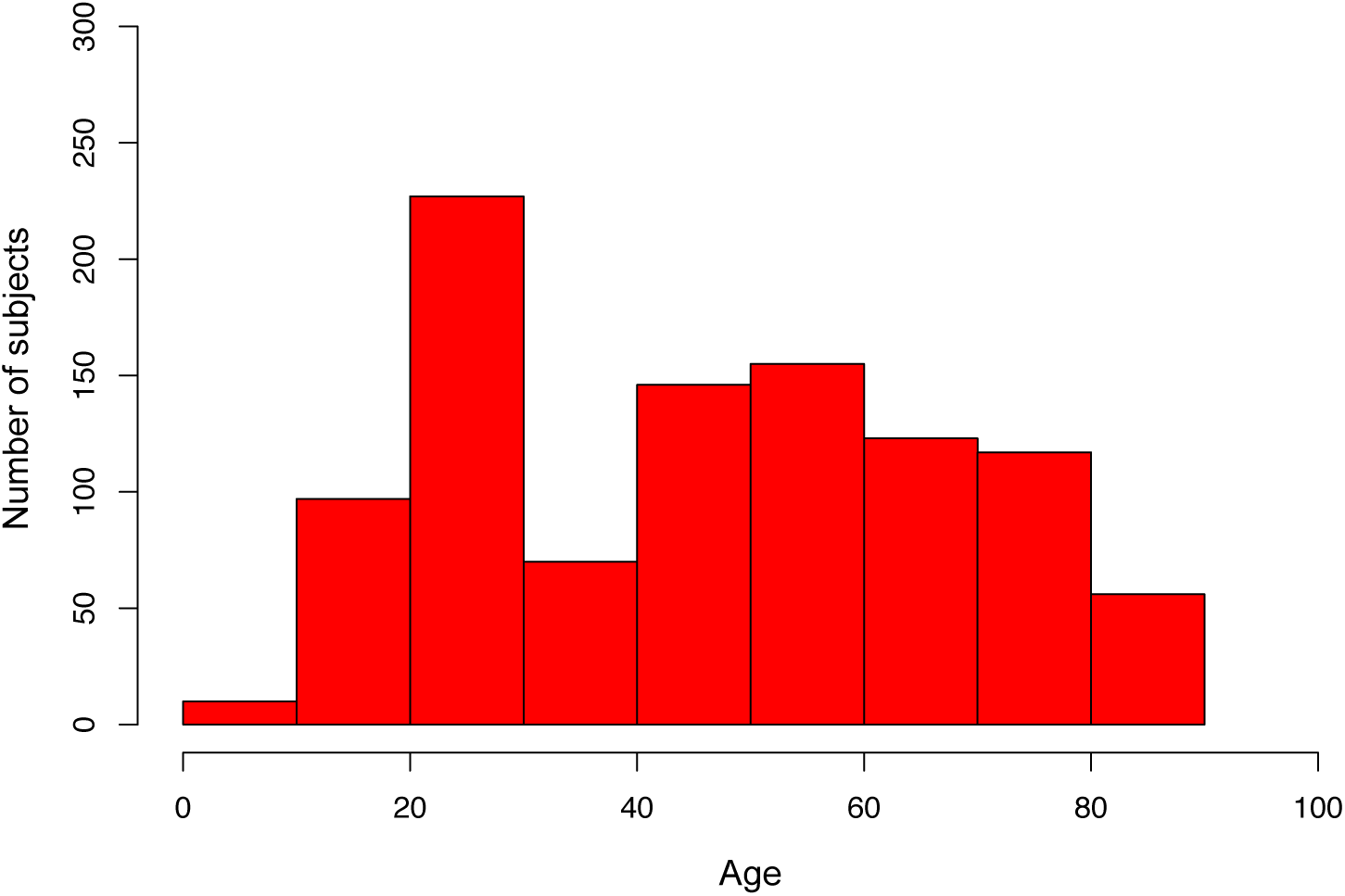
Age distribution of the Stanford 1KIP cohort.

**Supplementary Figure 3.**
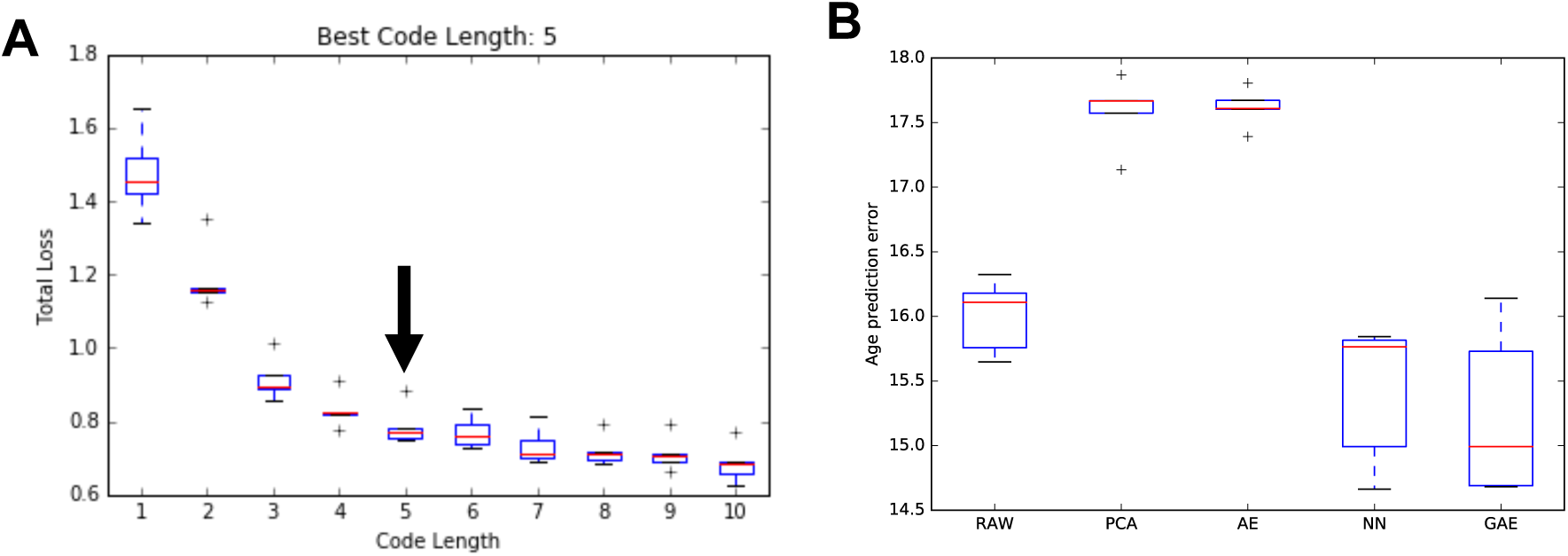
Estimation of the GAE code length and accuracy of age prediction. We used 5-fold cross-validation to identify the best code length, among lengths from 1 to 10. We selected the length of code k, whose performance was not statistically significantly worse than that of longer codes (paired t-test p-value > 0.05). Within each fold we performed nested 3-fold cross-validation to select hyper-parameters (depth, weight decay and guidance-ratio). In our experiment, the best code length is 5 (**A**) as adding one more code (6) does not significantly improve the total loss (p-value = 0.18). After obtaining the best code length as 5, we used the 5-fold-cross-validation to select the best hyper-parameter setting (depth = 2, guidance-ratio = 0.2, L2 = 0.001) on all GAE with code length 5. Finally, we trained the GAE on the whole dataset with the selected best hyper-parameter setting and obtained the predictive function as the inflammatory clock predictor. The GAE method outperforms linear methods for protein data reconstruction and prediction of chronological age (**B**).

**Supplementary Figure 4.**
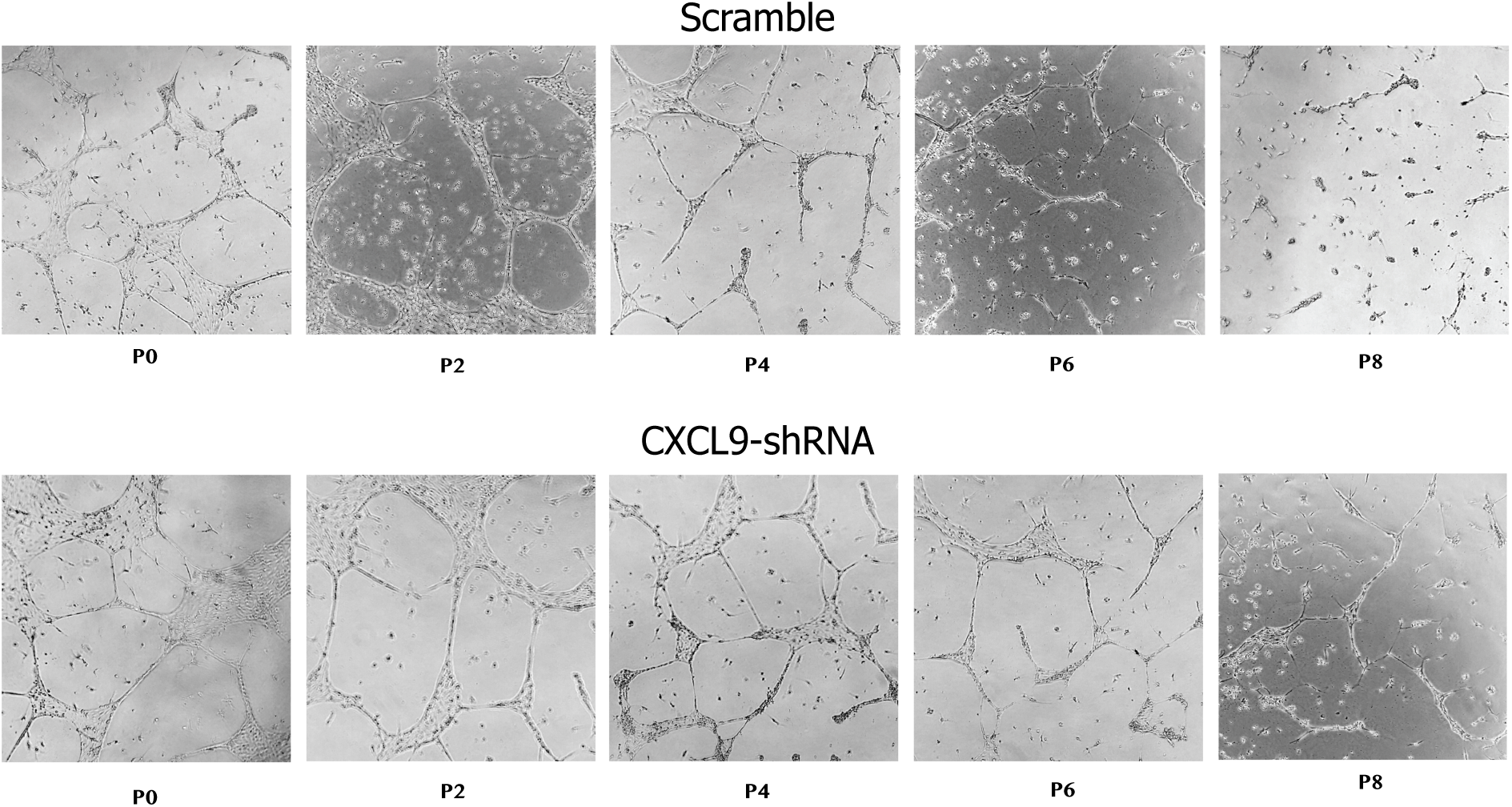
Validation of the effects of CXCL9 on endothelial function. Representative images of capillary-like networks from scramble- and CXCL9-KD hiPSC-ECs show that CXCL9-KD hiPSC-ECs retain their capacity to form tubes even at later passages when compared to scramble that showed impaired tube formation towards later passages of hiPSC ECs.

**Supplementary Figure 5.**
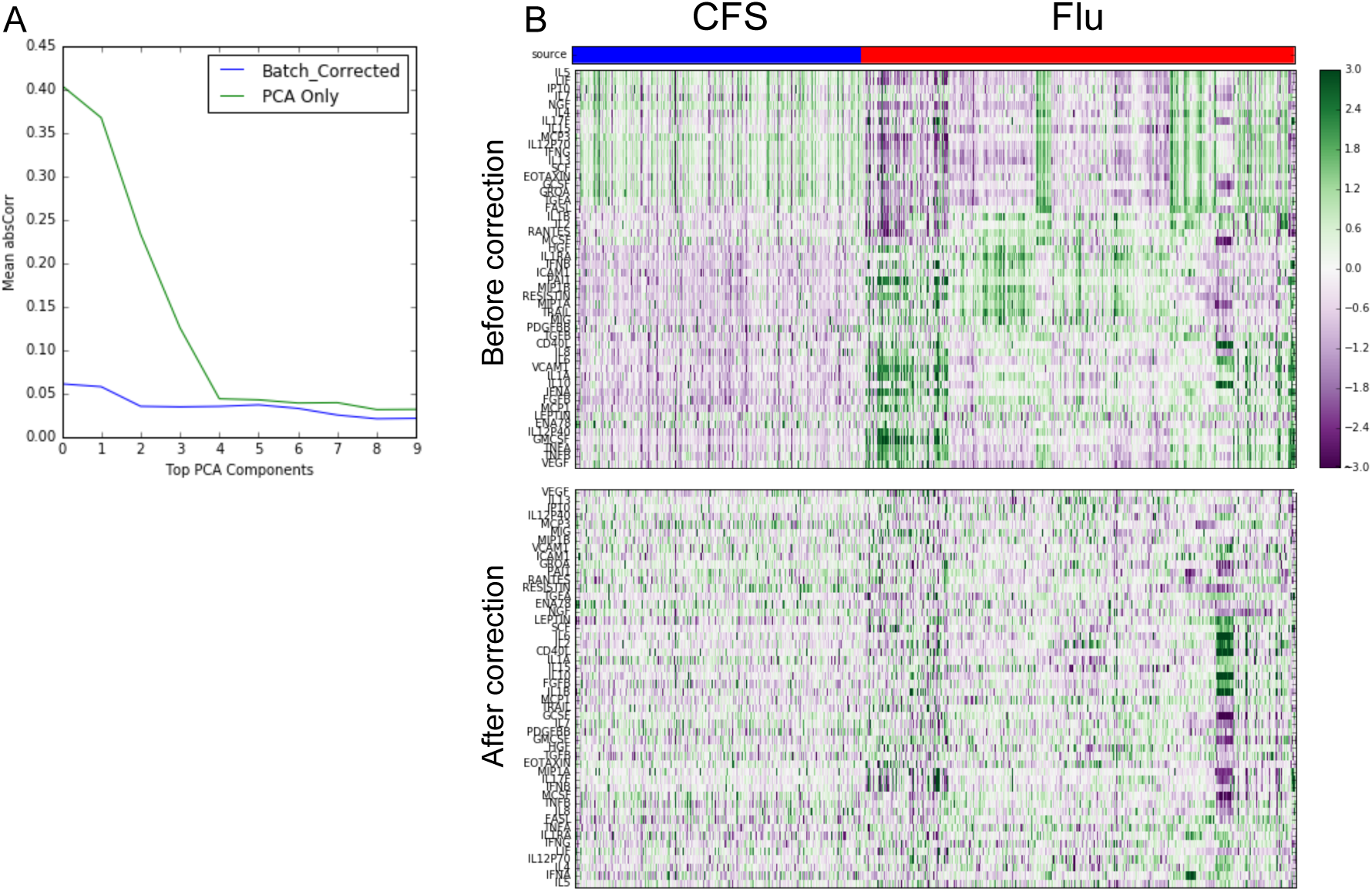
Elimination of batch effect for serum immune protein data. Immune protein data from serum samples were subjected to normalization and batch correction procedures (See Methods) to ensure data from different sources can be combined and used as a whole. (A) Spearman correlation between immune protein features and batch ID shows a strong dependency of data source on top 4 components (raw data, green line), which reaches a steady state after component 5. Data normalization and batch correction removes batch effect as indicated by lower mean absolute Spearman correlation between all features and batch id (blue line), which indicates impossibility to distinguish sample source from corrected data. (B) Upper panel: immune protein expression heatmap of uncorrected data, Lower panel: immune protein expression heatmap of corrected data.

**Supplementary Table 1.** Available data for the 1000 Immunomes Project.

|  | *# Features* | *Sample size* |
| --- | --- | --- |
| Cell subsets | **25** | **935** |
| Immune proteins | **50** | **1001** |
| Gene expression* | **6149** | **394** |
| Cell stimulations | **84** | **818** |
| Latent CMV | **1** | **748** |
| BMI | **1** | **724** |
| Clinical questionnaire | **57** | **902** |
| Cardiovascular phenotyping | **37** | **97** |

\*low variant genes removed

**Supplementary Table 2.**
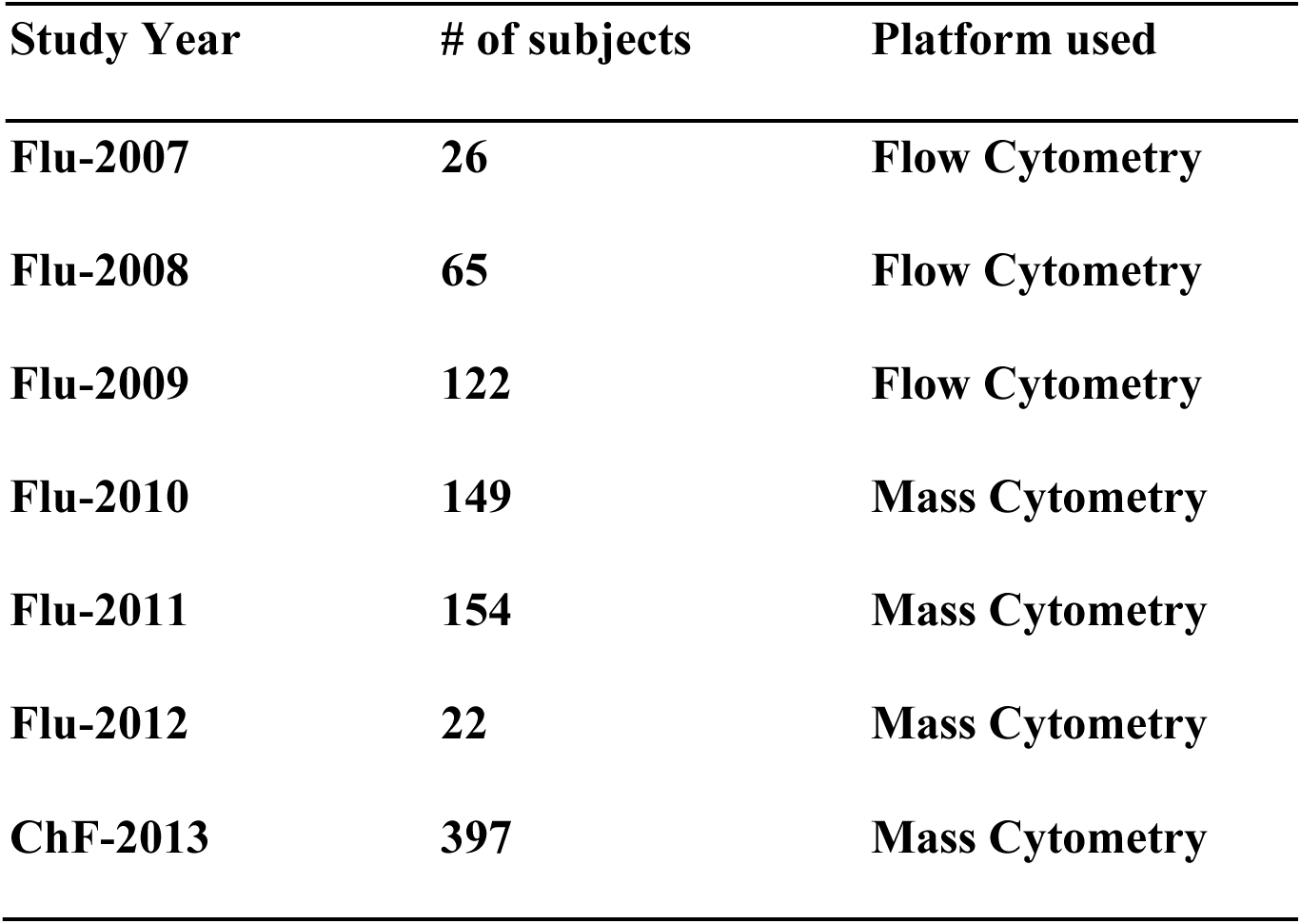
Number of subjects included, and analytical platform used for each study year.

| **Study Year** | **# of subjects** | **Platform used** |
| --- | --- | --- |
| **Flu-2007** | **26** | **Flow Cytometry** |
| **Flu-2008** | **65** | **Flow Cytometry** |
| **Flu-2009** | **122** | **Flow Cytometry** |
| **Flu-2010** | **149** | **Mass Cytometry** |
| **Flu-2011** | **154** | **Mass Cytometry** |
| **Flu-2012** | **22** | **Mass Cytometry** |
| **ChF-2013** | **397** | **Mass Cytometry** |

**Supplementary Table 3.**
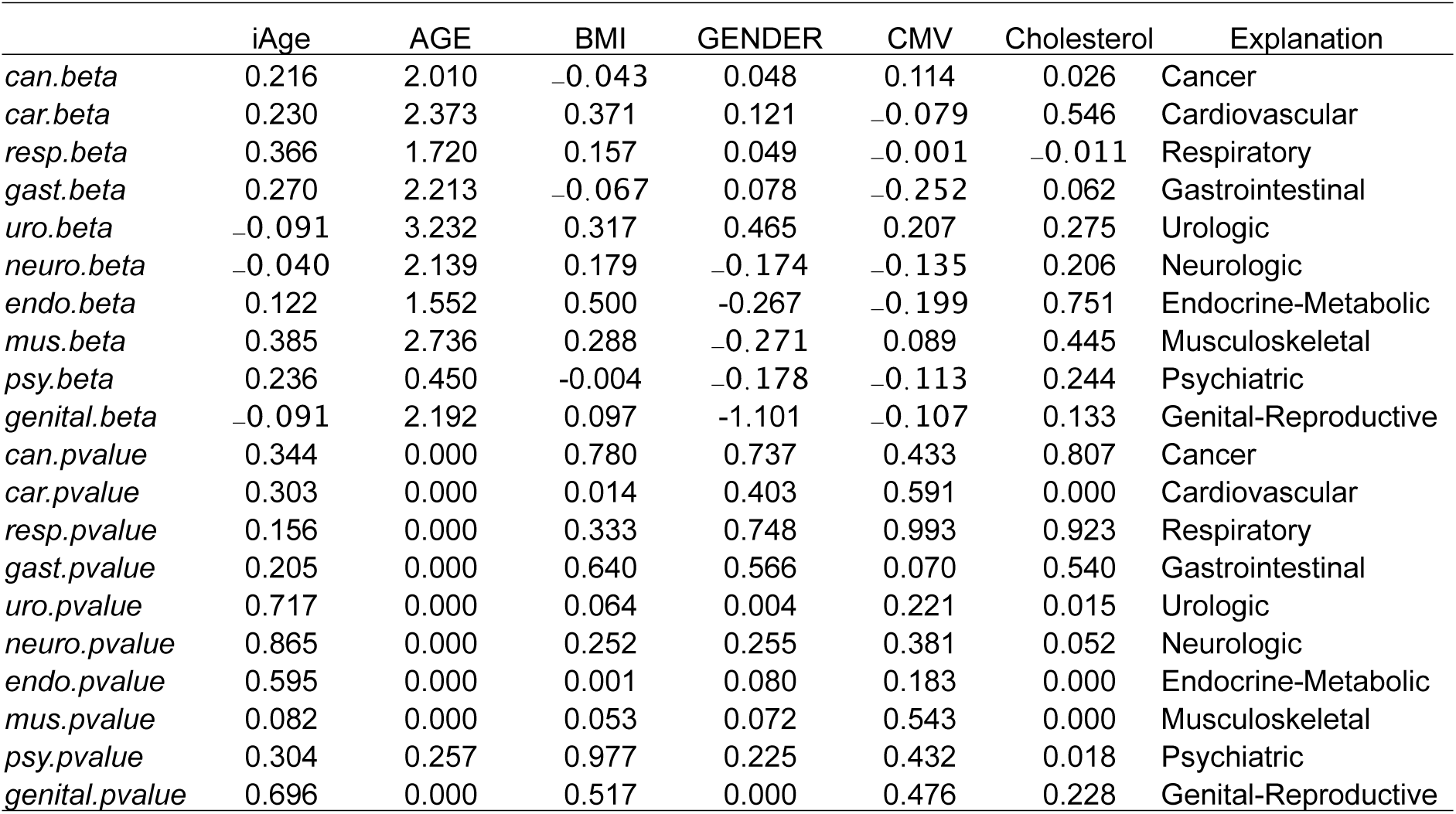
Regression coefficients and p-values for iAge and covariates in predicting individual disease items.

|  | iAge | AGE | BMI | GENDER | CMV | Cholesterol | Explanation |
| --- | --- | --- | --- | --- | --- | --- | --- |
| *can.beta* | 0.216 | 2.010 | -0.043 | 0.048 | 0.114 | 0.026 | Cancer |
| *car.beta* | 0.230 | 2.373 | 0.371 | 0.121 | -0.079 | 0.546 | Cardiovascular |
| *resp.beta* | 0.366 | 1.720 | 0.157 | 0.049 | -0.001 | -0.011 | Respiratory |
| *gast.beta* | 0.270 | 2.213 | -0.067 | 0.078 | -0.252 | 0.062 | Gastrointestinal |
| *uro.beta* | -0.091 | 3.232 | 0.317 | 0.465 | 0.207 | 0.275 | Urologic |
| *neuro.beta* | -0.040 | 2.139 | 0.179 | -0.174 | -0.135 | 0.206 | Neurologic |
| *endo.beta* | 0.122 | 1.552 | 0.500 | -0.267 | -0.199 | 0.751 | Endocrine-Metabolic |
| *mus.beta* | 0.385 | 2.736 | 0.288 | -0.271 | 0.089 | 0.445 | Musculoskeletal |
| *psy.beta* | 0.236 | 0.450 | -0.004 | -0.178 | -0.113 | 0.244 | Psychiatric |
| *genital.beta* | -0.091 | 2.192 | 0.097 | -1.101 | -0.107 | 0.133 | Genital-Reproductive |
| *can.pvalue* | 0.344 | 0.000 | 0.780 | 0.737 | 0.433 | 0.807 | Cancer |
| *car.pvalue* | 0.303 | 0.000 | 0.014 | 0.403 | 0.591 | 0.000 | Cardiovascular |
| *resp.pvalue* | 0.156 | 0.000 | 0.333 | 0.748 | 0.993 | 0.923 | Respiratory |
| *gast.pvalue* | 0.205 | 0.000 | 0.640 | 0.566 | 0.070 | 0.540 | Gastrointestinal |
| *uro.pvalue* | 0.717 | 0.000 | 0.064 | 0.004 | 0.221 | 0.015 | Urologic |
| *neuro.pvalue* | 0.865 | 0.000 | 0.252 | 0.255 | 0.381 | 0.052 | Neurologic |
| *endo.pvalue* | 0.595 | 0.000 | 0.001 | 0.080 | 0.183 | 0.000 | Endocrine-Metabolic |
| *mus.pvalue* | 0.082 | 0.000 | 0.053 | 0.072 | 0.543 | 0.000 | Musculoskeletal |
| *psy.pvalue* | 0.304 | 0.257 | 0.977 | 0.225 | 0.432 | 0.018 | Psychiatric |
| *genital.pvalue* | 0.696 | 0.000 | 0.517 | 0.000 | 0.476 | 0.228 | Genital-Reproductive |

**Supplementary Table 4.**
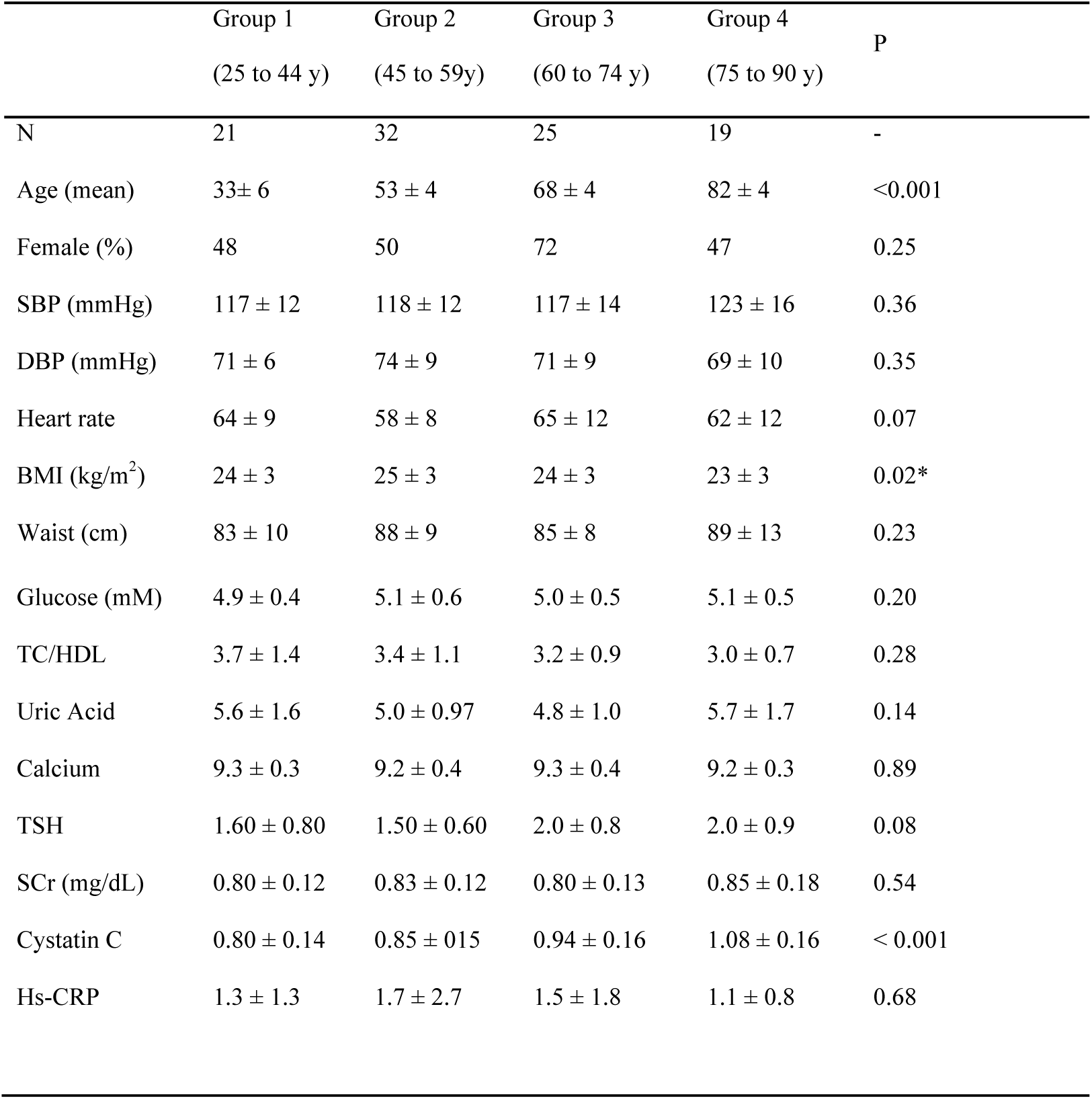
Validation study. Baseline clinical and demographic data.

|  | Group 1<br>(25 to 44 y) | Group 2<br>(45 to 59y) | Group 3<br>(60 to 74 y) | Group 4<br>(75 to 90 y) | P |
| --- | --- | --- | --- | --- | --- |
| N | 21 | 32 | 25 | 19 | - |
| Age (mean) | 33± 6 | 53 ± 4 | 68 ± 4 | 82 ± 4 | <0.001 |
| Female (%) | 48 | 50 | 72 | 47 | 0.25 |
| SBP (mmHg) | 117 ± 12 | 118 ± 12 | 117 ± 14 | 123 ± 16 | 0.36 |
| DBP (mmHg) | 71 ± 6 | 74 ± 9 | 71 ± 9 | 69 ± 10 | 0.35 |
| Heart rate | 64 ± 9 | 58 ± 8 | 65 ± 12 | 62 ± 12 | 0.07 |
| BMI (kg/m<sup>2</sup>) | 24 ± 3 | 25 ± 3 | 24 ± 3 | 23 ± 3 | 0.02* |
| Waist (cm) | 83 ± 10 | 88 ± 9 | 85 ± 8 | 89 ± 13 | 0.23 |
| Glucose (mM) | 4.9 ± 0.4 | 5.1 ± 0.6 | 5.0 ± 0.5 | 5.1 ± 0.5 | 0.20 |
| TC/HDL | 3.7 ± 1.4 | 3.4 ± 1.1 | 3.2 ± 0.9 | 3.0 ± 0.7 | 0.28 |
| Uric Acid | 5.6 ± 1.6 | 5.0 ± 0.97 | 4.8 ± 1.0 | 5.7 ± 1.7 | 0.14 |
| Calcium | 9.3 ± 0.3 | 9.2 ± 0.4 | 9.3 ± 0.4 | 9.2 ± 0.3 | 0.89 |
| TSH | 1.60 ± 0.80 | 1.50 ± 0.60 | 2.0 ± 0.8 | 2.0 ± 0.9 | 0.08 |
| SCr (mg/dL) | 0.80 ± 0.12 | 0.83 ± 0.12 | 0.80 ± 0.13 | 0.85 ± 0.18 | 0.54 |
| Cystatin C | 0.80 ± 0.14 | 0.85 ± 0.15 | 0.94 ± 0.16 | 1.08 ± 0.16 | < 0.001 |
| Hs-CRP | 1.3 ± 1.3 | 1.7 ± 2.7 | 1.5 ± 1.8 | 1.1 ± 0.8 | 0.68 |

